# Selective autophagy fine-tunes Stat92E activity by degrading Su(var)2-10/PIAS during glial injury signaling in *Drosophila*

**DOI:** 10.1101/2024.08.28.610109

**Authors:** Virág Vincze, Zsombor Esküdt, Erzsébet Fehér-Juhász, Aishwarya Sanjay Chhatre, András Jipa, Martin Csordós, Anna Rita Galambos, Dalma Feil-Börcsök, Gábor Juhász, Áron Szabó

## Abstract

Glial immunity plays a pivotal role in the maintenance of nervous system homeostasis and in responses to stress conditions, including neural injuries. The transcription factor Stat92E is activated independently of the canonical JAK/STAT pathway in *Drosophila* glial cells following central nervous system injury to shape glial reactivity towards degenerating axons. However, the upstream regulatory mechanisms governing Stat92E activation remain elusive. Here we reveal that selective autophagy mediates degradation of the PIAS SUMO ligase family member Stat92E repressor, Su(var)2-10 in glia. Autophagic elimination of Su(var)2-10 mediated by its co-localization and interaction with the core autophagy factor Atg8a is required for efficient Stat92E-dependent transcription after injury. In line with this, we demonstrate that autophagy is essential for the upregulation of an innate immune pathway in glial cells following axon injury, characterized by the induction of *virus-induced RNA 1* (*vir-1*). We propose that autophagic Su(var)2-10 breakdown controls Stat92E activation to allow glial reactivity. These findings identify a critical role for autophagy in glial immunity as part of neural injury responses.

## Introduction

The brain tissue in homeostatic conditions is constantly monitored by glia to mount an enhanced response in face of a challenge. Microglia persistently sample the environment: their elongated processes search for signals emitted from dying, damaged or infected cells. These trigger morphological and signaling changes in microglia to induce a transformation into a reactive state. Astrocytes undergo a similar transition upon stimuli. Reactive glia have distinct transcriptional signatures as exemplified by disease-associated microglia in neurodegeneration^1^. They sense damage- and pathogen-associated molecular patterns (DAMPs and PAMPs, respectively), aka. find-me and eat-me cues via dedicated receptors to mount an immune response. They also release various chemokines and cytokines to control neuroinflammation^2^. Upon central nervous system (CNS) injury, these receptors trigger signaling events that involve the MAPK, JAK-STAT and NF-κB pathways^2^. The transcriptomic changes in reactive glia show a major contribution from these signaling networks^3–7^.

The JAK-STAT pathway is an important but incompletely characterized arm of glial immunity, especially when compared to the NF-κB pathway^8^. Most interleukin and interferon receptors signal through JAK-STAT in mammals and induce the expression of either pro- or anti-inflammatory cytokines depending on the STAT paralog involved, both in peripheral immune cells and in glia^8,9^. In neurodegenerative diseases, the JAK-STAT pathway contributes to neuroinflammation. Increased interferon-dependent microglial Stat1 activation is observed in models of Alzheimer’s disease, causing complement-dependent synapse elimination^10^.

Research on *Drosophila* has elucidated key early glial responses to axon injury.^11–15^ The single STAT family transcription factor Stat92E upregulates the main glial phagocytic receptor *draper* (*drpr*) and matrix metalloproteinase *Mmp1* to induce glial reactivity^16,17^. Interestingly, this response does not rely on canonical pathway components such as JAK kinase (Hopscotch/Hop in flies) or its upstream receptor Domeless (Dome)^16^. Instead, the Drpr receptor and its downstream signaling partners involved in Rac1 activation are required for glial Stat92E activation after injury^16^. Insulin-like signaling also promotes glial Stat92E activity^18^. How Rac1 relays signaling to Stat92E is not known^19^. However, besides Stat92E, the transcription factor AP-1 also promotes *drpr* upregulation^12^ and Draper/Stat92E/AP-1 activity contributes to proper *Mmp1* expression after axon injury^17^.

Macroautophagy (simply referred to as autophagy) is a membrane-limited intracellular degradation pathway that removes cytoplasmic components including protein aggregates and organelles^19^. Upon induction by starvation or other stress conditions, an Atg1 (ULK1/2) complex initiates the formation of autophagic membranes. A network of Atg proteins mediate formation a cup-shaped phagophore that gives rise to an autophagosome, which transports cargo to lysosomes. This process requires Atg8a lipidation via two ubiquitin-like conjugation systems: Atg8a gets lipidated and thus membrane-bound by the Atg16/Atg5-Atg12 complex. Autophagosomes fuse with lysosomes where cargo breakdown and recycling take place.

A targeted variation on this theme is selective autophagy. During this process, cargo receptors recognize specific cellular components ranging from organelles to single molecules and recruit the autophagosome biogenesis machinery irrespective of nutritional status^20,21^. Cargo receptors can bind to ubiquitylated material or they can be embedded in target organelles, but labile proteins can also feature receptor-like motifs that mediates direct degradation^20,21^.

Since Stat92E activation is pivotal to glial adaptation to challenges, we need to gain a better insight into its regulatory mechanisms. Here we show that autophagy clears the direct Stat92E repressor PIAS/Su(var)2-10 in wing glia after wing transection, thereby gating Stat92E-dependent transcriptional regulation. Interestingly, *drpr* expression is not responsive to injury in these peripheral glial types. We find that another Stat92E target, *vir-1* is induced in an autophagy-dependent manner after wing transection.

## Results

### Selective autophagy is necessary for Stat92E but not for JNK pathway activation

To better understand nervous system adaptation after injury, we investigated whether autophagy impacts signaling pathways in glia. Two well-characterized immune pathways in glial responses are the JAK-STAT and JNK pathways. We used a well-established *Drosophila* wing injury model^22,23^ (Fig. S1A) to measure transgenic reporter upregulation for either of these pathways. As in the antennal lobe in the central nervous system (CNS) after antennal ablation^11,12,16^, both the Stat92E reporter *10xStat92E-GFP*^24^ and the AP-1 reporter *TRE-EGFP*^25^ showed robust activation after injury in the wing nerve (Fig. 1A,B, S1B,C). Their expression pattern coincided with a glially driven myr::tdTomato signal (Fig. S1B,C), similarly to the glial upregulation of Stat92E and AP-1 reporters in the CNS. In order to establish the dynamics of *10xStat92E-GFP* upregulation in the wing nerve, we first followed GFP reporter intensity at different time points after injury (Fig. S1D,E). *10xStat92E-GFP* signal intensity reached its peak 3 days post wing injury (3 dpi), so we measured reporter activation levels at this time point. We first depleted the core autophagy factor *Atg16* specifically in glia using *repo-Gal4*. While *Atg16* silencing did not affect *TRE-EGFP* expression, it abrogated *10xStat92E-GFP* activation after injury (Fig. 1A-C). We also measured Stat92E reporter activity in *Atg5 (Atg5^5cc5^)* and *Atg8a (Atg8a^Δ12^)* null mutant (Fig. S2A) backgrounds and observed a comparable effect to *Atg16* knockdown (Fig. 1D-F). Importantly, lack of STAT activation could be rescued by an endogenous promoter-driven *Atg5* transgene (*Atg5::3xHA*) in the *Atg5-*deficient background (Fig. 1D,F).

**Figure 1.**
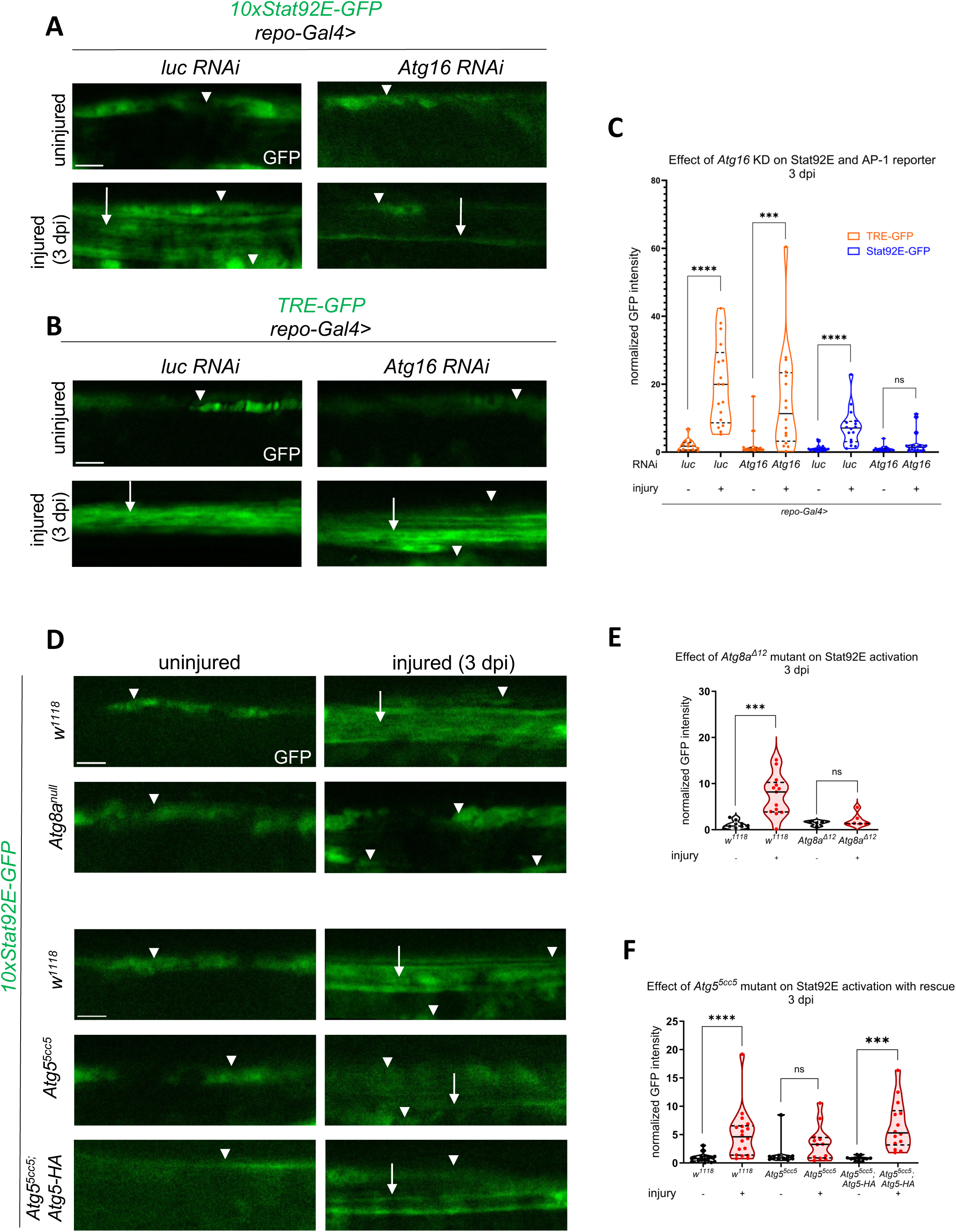
*Atg16* is required for Stat92E-, but not for AP-1-dependent transcription during glial reactivity. (A,B) Single-slice images showing the effect of *Atg16* and control (firefly luciferase: *luc*) knockdowns in glia on the *10xStat92E-GFP* and *TRE-EGFP* reporters in uninjured and injured wing nerves at 3 dpi (days post injury)(see also Fig. S1A). RNAi expression was induced by glia-specific *repo-Gal4*. (C) Quantitative analysis of the activity of the reporter constructs (A,B) based on normalized GFP intensity. Unpaired, two-tailed Mann-Whitney test was applied in each comparison. **** p < 0.0001, *** p = 0.0004, ns = 0.0743. n = 13, 17, 13, 16, 18, 16, 16, 17. (D) Single-slice images of wing nerves showing the *10xStat92E-GFP* signal at 3 dpi in *Atg8a^Δ12^* and *Atg5^5cc5^*mutant backgrounds. The *Atg5^5cc5^* phenotype can be rescued by introducing *Atg5::3xHA,* an endogeneous promoter-driven *Atg5* transgene. (E,F) Quantitative analysis of *10xStat92E-GFP* activation based on normalized GFP intensity. Unpaired, two-tailed *t*-test (*w^1118^(E), Atg5^5cc5^::3xHA*) and unpaired, two-tailed Mann–Whitney test (*w^1118^(F), Atg8a^Δ12^, Atg5*^5cc5^) were used for statistics. **** p < 0.0001, *** p = 0.0002, ns (E) > 0.9999, ns (F) = 0.1179. n (E) = 10, 13, 4, 6. n (F) = 17, 18, 11, 12, 11, 14. Truncated violin plots are shown with median and quartiles. Arrows point to glial reporter expression, while arrowheads denote the epithelial GFP signal surrounding the wing vein. Scale bar: 5 µm.

Atg5, Atg8a and Atg16 are not only involved in autophagy but also in autophagy-related processes including LC3-associated phagocytosis (LAP), which we have recently characterized in glia in relation to axon debris degradation^26^. Therefore, we set out to distinguish the contribution of these two processes to Stat92E signaling. Atg13 is a key subunit of the autophagy-initiating Atg1 kinase complex, which does not participate in LAP. *Atg13* silencing in glia also abrogated Stat92E reporter activation (Fig. 2A,C). Vice versa, Rubicon is dispensable for autophagy, but it is required for LC3-associated phagocytosis. Unlike *Atg13* knockdown, *Rubicon* RNA interference (RNAi) did not prevent Stat92E activation (Fig. 2B,D). We thus conclude that it is canonical autophagy and not LAP that sustains Stat92E signaling in glia.

**Figure 2.**
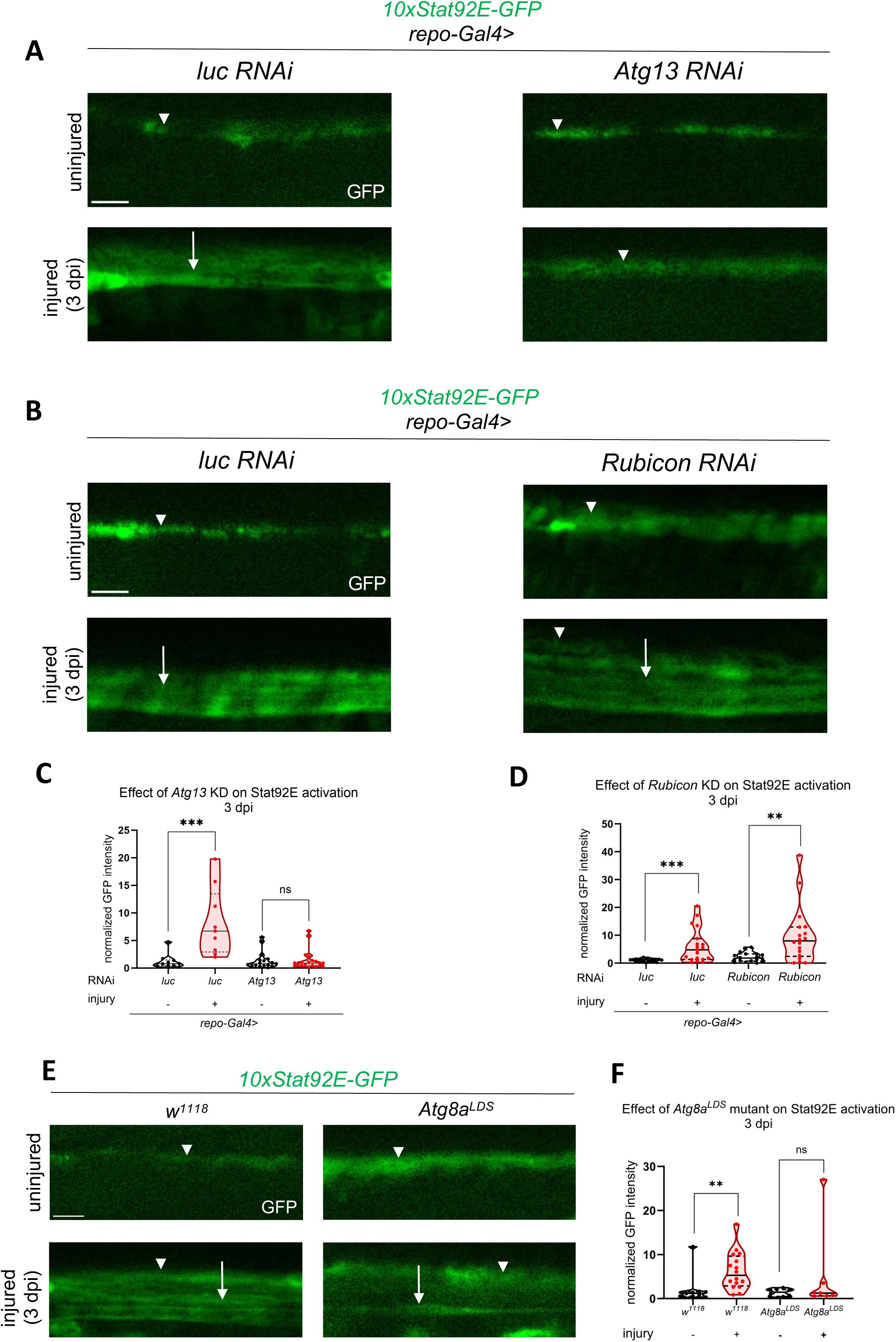
Selective autophagy drives Stat92E-dependent transcription in injury signaling. (A,B) Single-slice images of wing nerves showing *10xStat92E-GFP* signal in different glial RNAi backgrounds. *Atg13* is required only for autophagy, while *Rubicon* is indispensable for LAP (LC3/Atg8a-associated phagocytosis). RNAi expression was induced by *repo-Gal4*. (C,D) Quantitative analysis of *10xStat92E-GFP* activation (A, B) based on normalized GFP intensity. Unpaired, two-tailed Mann–Whitney test was used for statistics. *** p(C) = 0.0002, *** p(D) = 0.0004, ** p = 0.0021, ns = 0.9195. n (C) = 10, 9, 18, 20. n (D)= 19, 19, 18, 18. (E) Single optical slices showing the effect of *Atg8a^LDS^* mutation on *10xStat92E-GFP* activation*. Atg8a^LDS^* has a mutation in its LIR docking site (LDS), which specifically eliminates LIR-dependent selective autophagy. (F) Quantitative analysis of *10xStat92E-GFP* activity (E) based on normalized GFP intensity. Unpaired, two-tailed Mann-Whitney test was used for each comparison. ** p = 0.0053, ns = 0.7789. n = 9, 16, 8, 7. Truncated violin plots are shown with median and quartiles. Arrows point to glial GFP, while arrowheads denote the epithelial GFP signal. Scale bar: 5 µm.

We aimed to further understand how autophagy specifically impacts Stat92E activation. Selective autophagy receptors (SARs) use their LC3-interacting region (LIR) to bind the LIR motif docking site (LDS) on the core autophagy protein Atg8a. SARs either directly interact with to-be-degraded cargos or with polyubiquitin (Ub) chains on cargos. LIRs can be also present in cargos themselves to facilitate their degradation via Atg8a binding. We thus evaluated a point mutant form of Atg8a that specifically disrupts the LDS to see if it influences Stat92E-dependent gene activation^27^. The homozygous *Atg8a^K48A/Y49A^* (*Atg8a^LDS^)* mutation indeed prevented Stat92E reporter upregulation following injury (Fig. 2E,F). Although silencing *ref(2)p/p62*, the only known Ub chain-binding SAR in *Drosophila* also showed a mild decrease in GFP reporter intensity, it did not have a statistically significant effect on *10xStat92E-GFP* activation (Fig. S2B,C). Of note, bulk autophagic degradation in glia did not change in response to injury based on the tandem tagged autophagic flux reporter mCherry-GFP-Atg8a expressed in wing glia (Fig. S2D)^28^. These data suggest that a negative regulator of Stat92E signaling is removed by selective autophagy, involving SAR(s) different from/in addition to Ref(2)P/p62.

### *vir-1* but not *drpr* is upregulated by autophagy-activated Stat92E signaling in glia after wing injury

What are the relevant targets of autophagy-regulated Stat92E activation in glial injury responses? It has been shown that following antennal ablation, baseline *drpr* expression in the brain increases via Stat92E-dependent transcription in response to Drpr receptor signaling, creating a positive feedback loop^16^. This is required for the efficient degradation of axon debris generated after injury. We monitored *drpr* transcription after neural injury to see whether it is under the influence of autophagic regulation. For this, we converted a MiMIC transposon insertion in *drpr* (*drpr*^MI07659^) into a *Trojan-Gal4*, in which Gal4 expression reflects *drpr* transcription (Fig. S3A). We expressed the recently developed transcriptional reporter *UAS-TransTimer*^29^ with our *drpr-Gal4.* TransTimer allows measuring the ratio of destabilized nuclear GFP versus a stable nuclear RFP to monitor dynamic changes in transcription. Surprisingly, *drpr-Gal4* flies did not show *TransTimer* upregulation after wing injury (Fig. S3B,C). To further confirm this unexpected result, we next drove *UAS-TransTimer* with an intronic *drpr* enhancer*-Gal4* (*dee7-Gal4*) that harbors a functional Stat92E binding site (Fig. S3A) and has been shown to upregulate transcription after injury^16^. First, we verified that *dee7-Gal4* expression is induced in the brain after antennal injury as described previously (Fig. S4A)^16^. However, we still failed to observe an increase in the number of RFP^+^ puncta 3 days after wing injury in *dee7-Gal4>TransTimer* flies (Fig. S4B,C). We thus conclude that glial *drpr* is not upregulated by wing injury and Drpr accumulation is apparently not a prerequisite for efficient debris processing within the wing nerve.

JAK-STAT signaling underlies many immune pathways including cytokine signaling and defense against viruses^30^. We screened immunity-related gene reporters (Supplementary Table 1) and found that *vir-1-GFP*^31^ (Fig. 3A) is robustly upregulated after wing injury (Fig. 3B,C). GFP expression under the control of the *vir-1* promoter overlapped with glially-driven myr::tdTomato, which clearly indicates *vir-1* induction in glia (Fig. S5A). Stat92E signaling-driven *vir-1* is one of the most strongly induced transcripts after viral infection in *Drosophila*^31^. Curiously, *vir-1* is markedly upregulated in glia specifically one day after traumatic brain injury but not later^32^. Brains one day post antennal ablation indeed showed a significant increase in *vir-1* reporter GFP levels in glia in and around the antennal lobe (Fig. 3D,E). *vir-1* has seven transcript isoforms*. RE* and *RD* are shorter and encompass only the 3’ half of the longer isoforms (*RA, RB, RG*). Therefore, their transcription initiation is offset relative to these longer variants. We found that after wing injury, *vir-1* isoforms *RA* and *RB* are upregulated while *RE* is not (Fig. S5B). This indicates that an injury-responsive enhancer resides in the proximity of the long isoform’s transcriptional start site.

**Figure 3.**
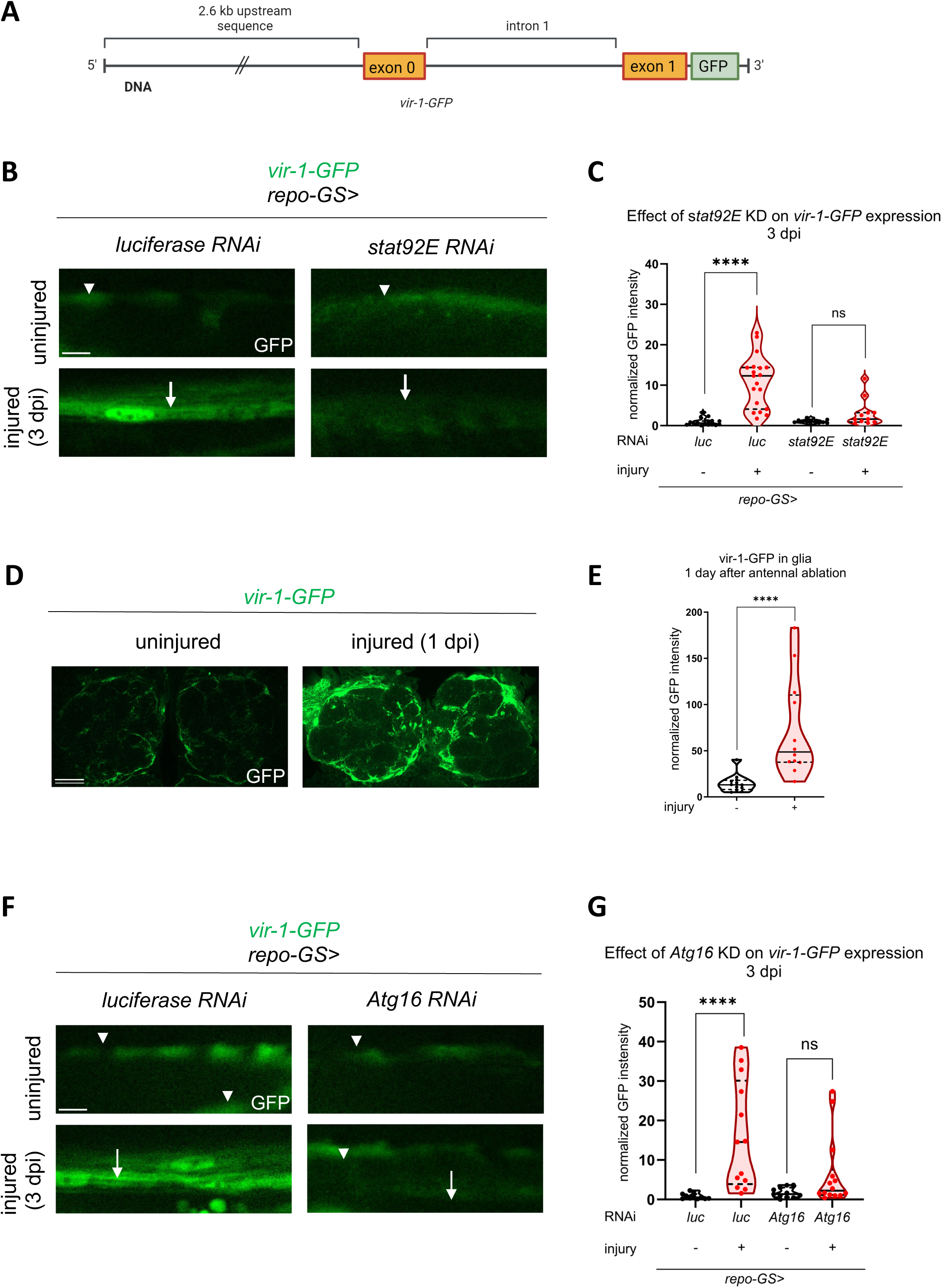
*vir-1* expression is regulated by Stat92E and is controlled by autophagy in glia after wing injury. (A) Schematic representation of the *vir-1-GFP* reporter construct: 2.6 kb of *vir-1* upstream sequences drive GFP expression^31^. The *vir-1* gene fragment includes the first intron and the first exon. (B) Stat92E-dependent activation of *vir-1-GFP* in glia after axonal injury. Single-slice images of wing nerves showing the *vir-1-GFP* signal upon glia-specific *stat92E* knockdown. Scale bar: 5 µm. (C) Quantification of *vir-1-GFP* signal shown in (B). Unpaired, two-tailed Mann–Whitney test was used for each pairwise comparison. **** p < 0.0001, ns = 0.0512. n = 16, 19, 12, 11. (D) *vir-1* induction in the brain after injury. Single-slice images showing the *vir-1-GFP* signal around the AL, in uninjured animals and 1 day after antennal ablation. Scale bar: 20 um. (E) Quantitative analysis of normalized GFP signal in the brain shown in (D). Data were quantified by using unpaired, two-tailed Mann–Whitney test. **** p < 0.0001. n = 12, 12. (F). *vir-1-GFP* induction upon injury in *Atg16* knockdown flies. Single-slice images of wing nerves showing the *vir-1-GFP* signal upon glial *Atg16* RNAi. Scale bar: 5 µm. (G) Unpaired, two-tailed Mann–Whitney test was used for both pairwise comparisons. **** p < 0.0001, ns = 0.1201. n = 11, 13, 11, 14. RNAi expression was induced by RU486 for 5-7 days after eclosion prior to injury and during the experiment. Arrows point to reporter expression in glia, while arrowheads denote the epithelium. Truncated violin plots are shown with median and quartiles.

To corroborate that *vir-1* is a glial Stat92E target, we silenced *stat92e* acutely in adult glia using *repo-GeneSwitch (repo-GS)* where RNAi expression was induced only in adults to circumvent potential developmental defects. It also enabled us to reduce *stat92e* levels in a narrow temporal window starting shortly before injury. *stat92e* knockdown abrogated *vir-1-GFP* induction in the wing nerve in response to injury (Fig. 3B,C). This prompted us to investigate whether autophagy impacts *vir-1* expression, similar to the *10xStat92E-GFP* reporter. We expressed *Atg16* RNAi in glia and followed *vir-1* induction after injury. While *vir-1* still tended to show some upregulation when *Atg16* was depleted in glia, this change was not statistically significant when compared to uninjured wings (Fig. 3F,G). Accordingly, we propose that autophagy contributes to glial reactive states at least in part by acting on this important innate immune locus via the regulation of Stat92E.

### Su(var)2-10 depletion enhances Stat92E activation in glia after wing injury

Next, we screened Stat92E regulators that may be responsible for Stat92E repression in glia. Since the upstream components of the JAK-STAT pathway, Hop and Dome are not involved in glial injury responses in flies^16^, we focused on interactors of Stat92E itself^33^. A direct repressor of Stat92E is Su(var)2-10 / Protein inhibitor of activated STAT (PIAS)^34^, a SUMO ligase whose mammalian orthologs PIAS1-4 SUMOylate and thereby inactivate STAT proteins^35^. We drove two independent *Su(var)2-10* RNAi constructs in adult glia using *repo-GS.* Strikingly, *Su(var)2-10* knockdown relieved Stat92E reporter repression predominantly following injury and not much in uninjured conditions (Fig. 4A-C). This raises the possibility that Su(var)2-10 elimination is important for efficient Stat92E-dependent activation after injury.

**Figure 4.**
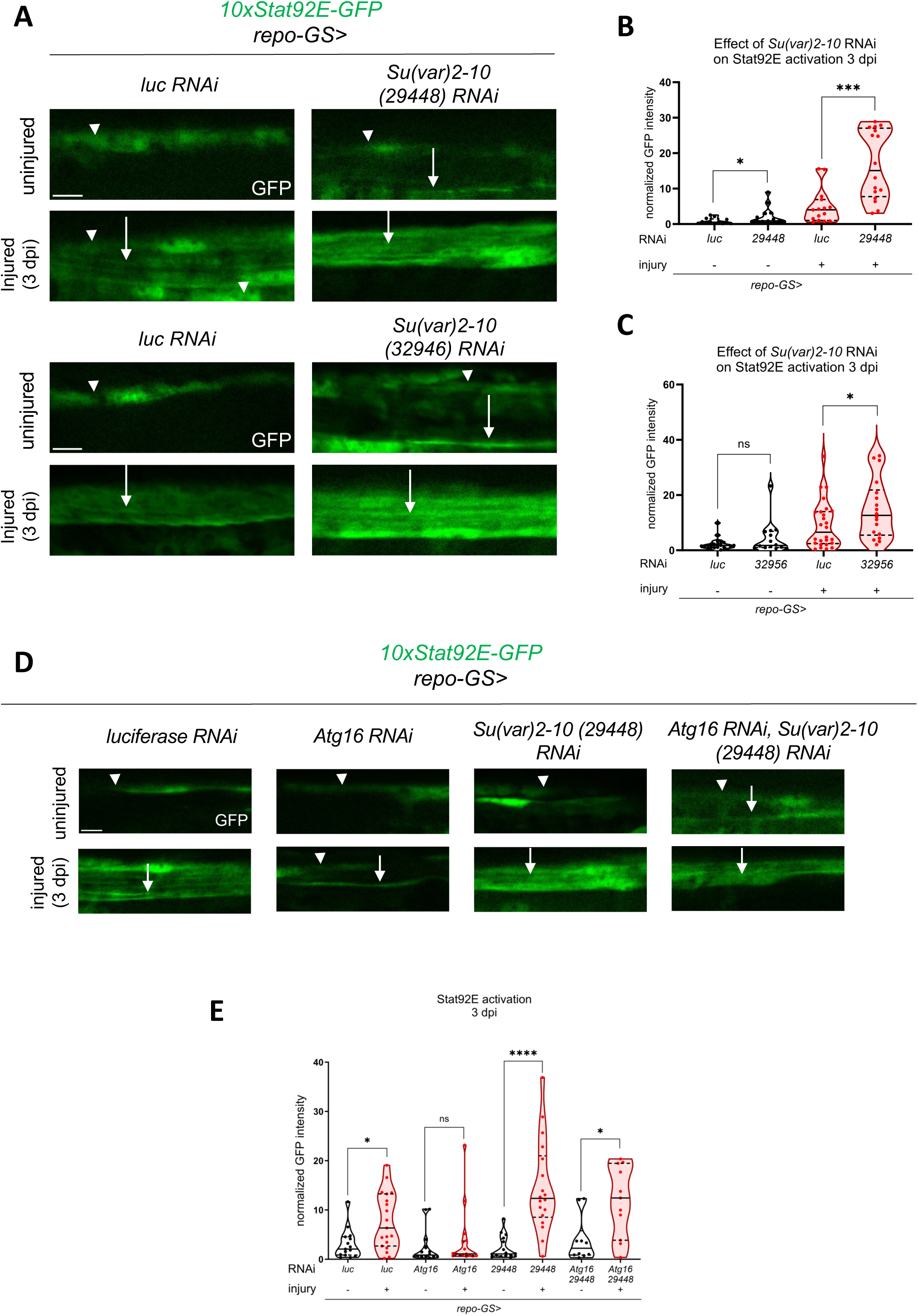
Su(var)2-10 is a Stat92E repressor in glia and mediates the autophagic modulation of Stat92E activity upon injury. (A) Single optical slices showing the disinhibition effect of two independent *Su(var)2-10* RNAi constructs on Stat92E activity (*10xSTAT92E-GFP*) using the *repo-GS* system in glia (B,C). Quantitative analysis of normalized *10xStat92E-GFP* signal shown in (A). Unpaired, two-tailed Mann-Whitney test was used for pairwise comparisons. *** p = 0.0002, * p (B) = 0.0120, * p (C) = 0.0482, ns = 0.4742. n (B) = 13, 13, 15, 16. n (C) = 21, 14, 26, 18. (D) Epistatic interaction between *Su(var)2-10* and *Atg16*. *Atg16* silencing impairs the activation of Stat92E after injury that is restored by simultaneous *Atg16* RNAi and *Su(var)2-10* knockdown. (E) Quantitative analysis of normalized *10xStat92E-GFP* signal shown in (D). Unpaired two-tailed Mann– Whitney test was used for all comparisons. **** p < 0.0001, * p (*luc*) = 0.0124, * p (*Atg16 + 29448*) = 0.0127, ns = 0.2939. n = 15, 19, 14, 16, 16, 18, 10, 11. RNAi expression was induced by RU486 feeding for 5-7 days after eclosion, prior to injury and during the experiment. Arrows point to glial GFP, while arrowheads denote the epithelial GFP signal. Scale bar: 5 µm. Truncated violin plots are shown with median and quartiles.

*Su(var)2-10* depletion enhanced glial Stat92E activity as opposed to the effect caused by the knockdown or abrogation of an autophagy-related component. We wondered whether Su(var)2-10 relays the repressive effect of autophagy disruption onto Stat92E reporter activation. We addressed this question via a genetic epistasis experiment. While knockdown of *Atg16* abrogated *10xStat92E-GFP* activation, depletion of *Su(var)2-10* in this background restored the normal magnitude of Stat92E response to injury (Fig. 4D,E, see also Fig. S6A,B for additional controls). This points to Su(var)2-10 as an important mediator of the autophagic regulation of Stat92E-dependent transcription.

### Autophagic degradation of Su(var)2-10 activates Stat92E upon injury

We further tested the hypothesis that autophagic degradation of Su(var)2-10 leads to Stat92E activation after injury. We depleted *Atg16* in glia and examined Su(var)2-10 levels in the adult brain following antennal ablation. Removal of antennae results in axon fragmentation in the antennal lobe that triggers reactivity in surrounding glia^15^. Su(var)2-10 is mainly nuclear due to its transcriptional repressor and chromatin organizer functions^34,36^. We expressed nuclear GFP in glia and inspected the co-localizing Su(var)2-10 pool. Su(var)2-10 levels were elevated in glia nuclei around the antennal lobe upon *Atg16* silencing in both uninjured and injured conditions (Fig. 5A-C). This was further validated by examining Su(var)2-10 abundance after antennal ablation in an *Atg101* null mutant in which autophagy initiation is defective (Fig. 5D-F). Interestingly, the accumulation of Su(var)2-10 was highly significant in injured brains, while the effect was much milder in uninjured brain glia of *Atg101* mutant animals. We confirmed the specificity of the Su(var)2-10 antibody on Western blots of Su(var)2-10::EGFP ectopically expressed in Schneider 2 R+ (S2R+) *Drosophila* cell culture (Fig. S6C) and homozygous *Su(var)2-10^MI03442^*loss-of-function mutant larval extracts (Fig. S6D). These results suggest that Su(var)2-10 is an autophagic cargo.

**Figure 5.**
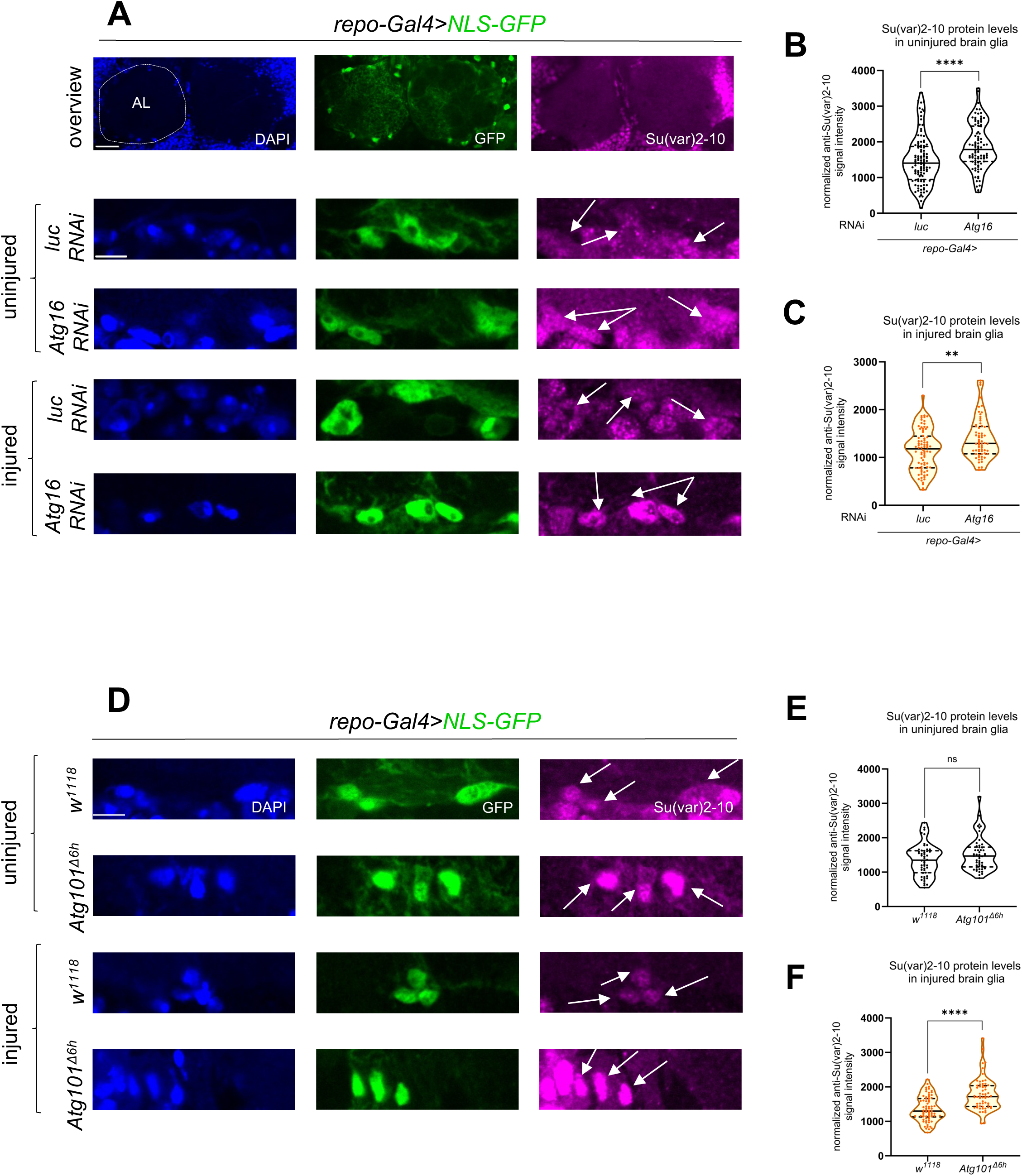
Su(var)2-10 levels are elevated in brain glia upon autophagy inhibition. (A,D) Single optical slices showing Su(var)2-10 abundance in brain glia around the antennal lobe (AL) in glial *luc* and *Atg16* KD (A) and *w^1118^* and *Atg101^Δ6h^* mutant (D) background, without injury and 1 day after antennal ablation, respectively. Su(var)2-10 is predominantly nuclear. Glial nuclei are labelled by *repo-Gal4>NLS-GFP.* Arrows denote glial nuclei. Note the accumulation of Su(var)2-10 in glia nuclei, and that a fraction of NLS-GFP is also present in the cytoplasm. Scale bars: 20 µm and 5 µm. (B,C) Quantitative analysis of Su(var)2-10 nuclear intensity normalized to background signal in single optical slices shown in (A). Unpaired, two-tailed Mann–Whitney test was used for statistics. **** p < 0.0001, ** p = 0.0033. n (B) = 109, 90. n (C) = 80, 64. (E, F) Quantitative analysis of Su(var)2-10 nuclear intensity normalized to background signal in single optical slices shown in (D). Unpaired, two-tailed Mann–Whitney test was used for statistics. **** p < 0.0001, ns = 0.1289. n (E) = 50, 50. n (F) = 70, 60. Truncated violin plots are shown with median and quartiles.

We noticed that nuclear Su(var)2-10 often colocalizes with glially expressed Atg8a in a punctate pattern in brains (Fig. 6A), raising the possibility that these proteins interact with each other. Although we did not find a predicted consensus LIR motif in Su(var)2-10, its interaction with Atg8a might be mediated by other motifs^21^. Su(var)2-10 is known to autoSUMOylate itself^37,38^. We found that SUMO colocalizes with structures that also contain Su(var)2-10 and Atg8a (Fig. 6A). Su(var)2-10 degradation must occur in the cytoplasm if it is an autophagy cargo^21^. Indeed, cytoplasmic structures double positive for Su(var)2-10 and Atg8a were readily detected in antennal lobe glia (Fig. 6B).

**Figure 6.**
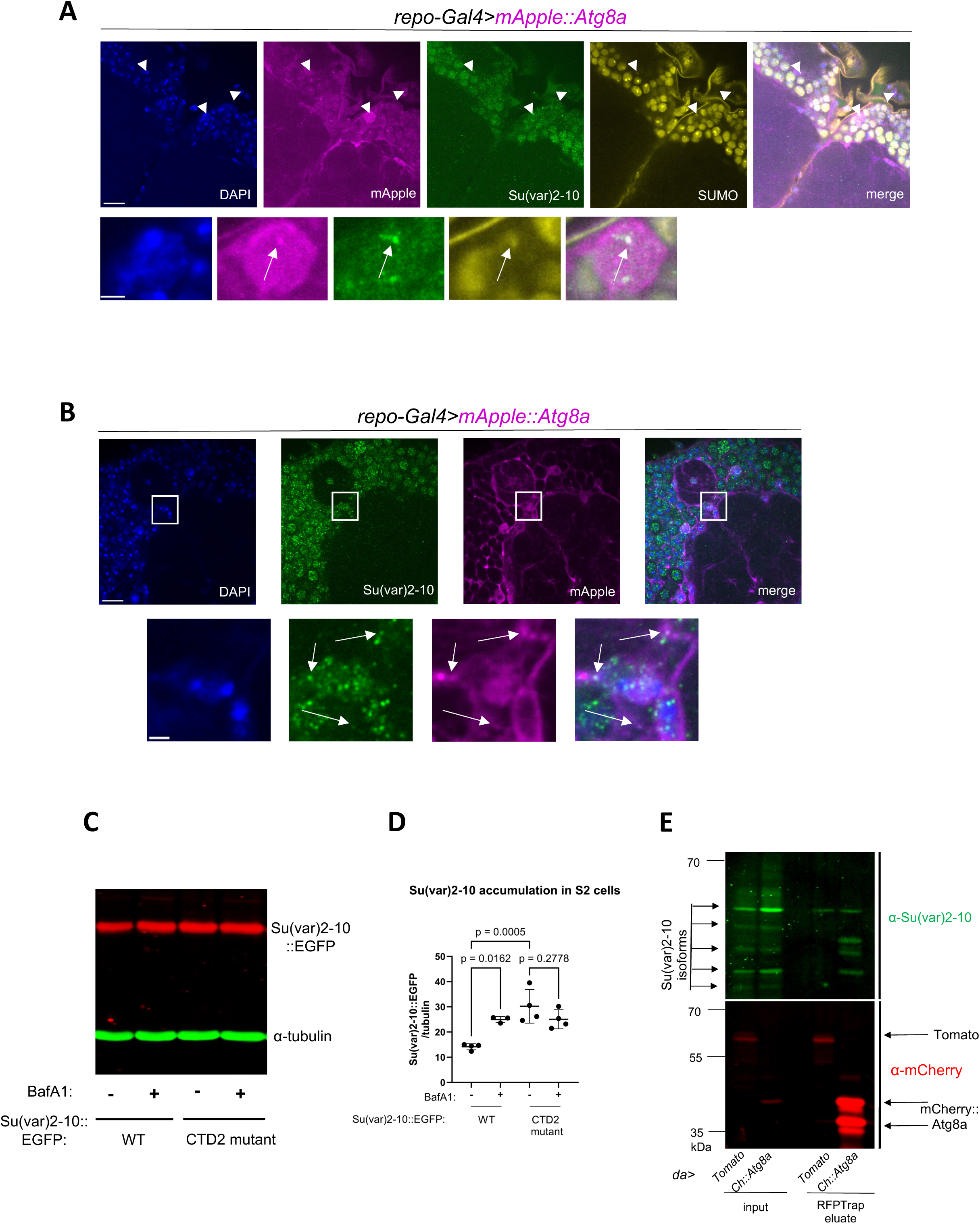
Su(var)2-10 is degraded by autophagy. (A) Single optical brain slice around the dorsal antennal lobe area is shown. Magnified images of the nuclei of interest are shown below. Arrowheads point to the glial nuclei of interest, while arrows denote the exact location of Su(var)2-10 - SUMO - Atg8a colocaliziation. Nuclei are stained with DAPI, native *mApple::Atg8a* signal from expression in glia by *repo-Gal4*, combined with anti-Su(var)2-10 and anti-SUMO immunostaining. Scale bar for brain images: 10 µm, scale bar for nucleus images: 2 µm. (B) Confocal single-slice multi-channel images show Su(var)2-10 - Atg8a colocalization in the cytoplasm of glial cells. Magnified images of the areas outlined by the rectangles can be seen below. Arrows point to the exact location of the colocalization. Nuclei are stained with DAPI, *mApple::Atg8a* expression is driven in glia by *repo-Gal4*, signal amplified by anti-mCherry staining, jointly with anti-Su(var)2-10 immunostaining. Scale bar for brain images: 10 µm, scale bar for nucleus images: 2 µm. (C) S2 cells transiently expressing wild-type and autoSUMOylation defective CTD2 mutant Su(var)2- 10::EGFP were treated with Bafilomycin A1 (BafA1) or the solvent DMSO for 4 hours, and total Su(var)2-10 levels were assessed by anti-GFP Western blot. α-tubulin serves as a loading control. (D) Quantification of Su(var)2-10 abundance normalized to α-tubulin as in (C). Shown is the mean and standard deviation, one-way ANOVA with Holm-Šídák test for multiple comparison correction was used for statistics. n = 4, 3, 4, 4. (E) Co-immunoprecipitation of Su(var)2-10 with mCherry::Atg8a. Ubiquitious *daughterless* (*da*)-*Gal4-*driven myr::tdTomato (Tomato) and mCherry::Atg8a (Ch::Atg8a) whole adult fly extracts were precipitated with RFPTrap beads. myr::tdTomato serves as a negative control. Eluates are shown on the right, crude extract inputs (10 ug) on the left. The two major bands in mCherry::Atg8a correspond to the lipidated and unlipidated forms. Western blots were probed by anti-mCherry and anti-Su(var)2-10.

**Fig. 7.**
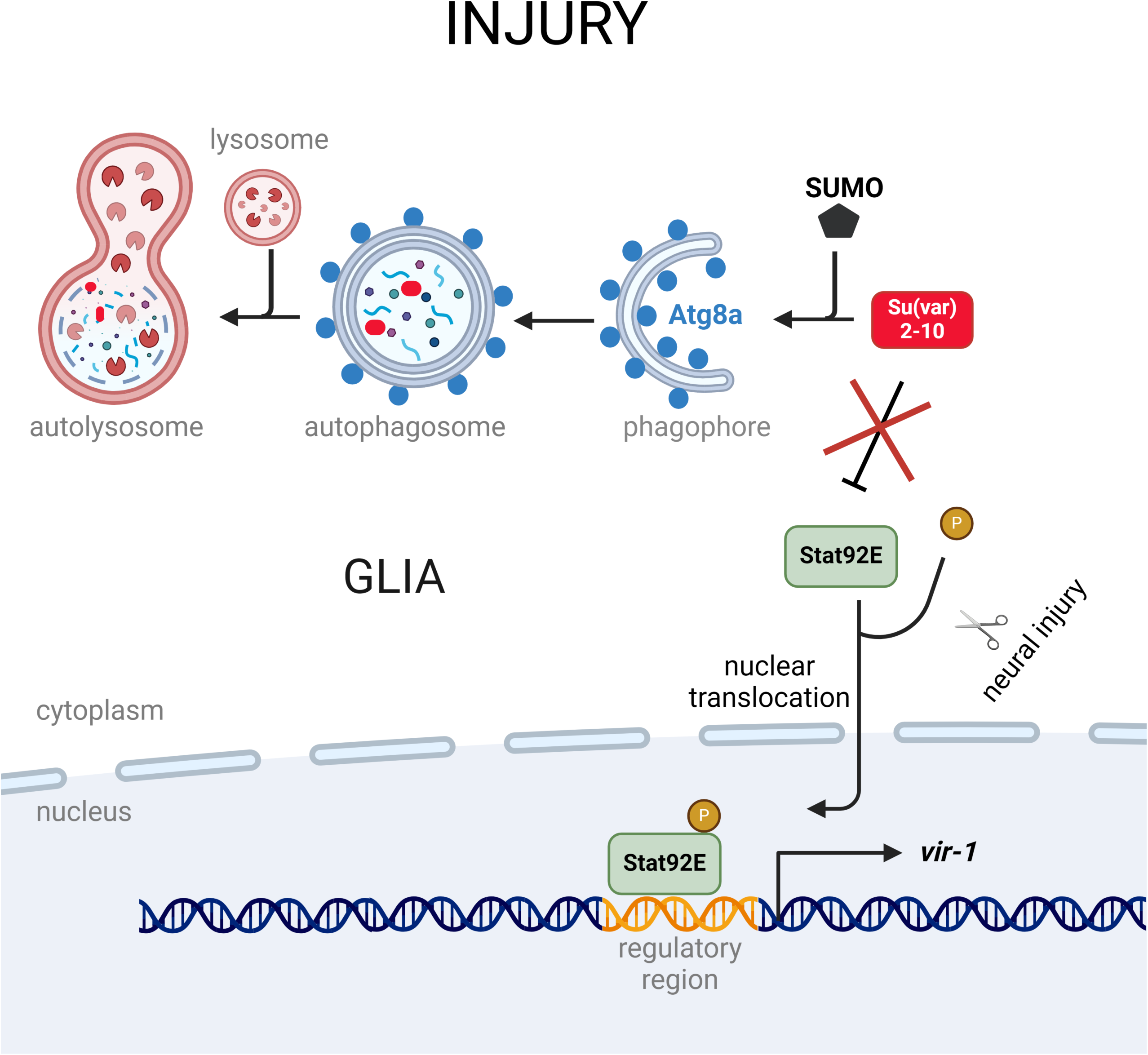
Schematic illustration of the autophagy-regulated Su(var)2-10/Stat92E axis in glia after neural injury. Upon nervous system injury, Stat92E is activated in glia by simultaneous phosphorylation and derepression. Su(var)2-10-mediated repression of Stat92E is relieved by selective autophagic degradation via the interaction of Atg8a and Su(var)2-10. Degradation of Su(var)2-10 involves autoSUMOylation. Stat92E phosphorylation concurrently occurs in response to injury. Activated Stat92E translocates to the nucleus and binds to Stat92E binding motifs in the regulatory region of genes that are induced after axon injury, such as *vir-1,* to mediate their transactivation.

To gain more insight into the degradation mechanism, we next tested Su(var)2-10 degradation in a heterologous system by transfecting cultured *Drosophila* S2R+ cells with *Su(var)2-10::EGFP* in the presence or absence of Bafilomycin A1 (BafA1) that inhibits both autophagosome fusion and lysosomal acidification. Su(var)2-10 levels increased upon BafA1 treatment, which indicates the lysosomal degradation of this protein (Fig. 6C,D). Is autoSUMOylation of Su(var)2-10 the lysosomal degradation signal? The C-terminal domain (CTD) of Su(var)2-10 interacts with SUMO. Mutating two residues in the CTD2 region disrupts SUMO binding and the autoSUMOylation of Su(var)2-10^38^. This leads to a decreased nuclear versus cytoplasmic Su(var)2-10 ratio *in vivo*. Using this construct, we found that baseline levels of CTD2 mutant form are increased relative to the wild-type protein, hinting to a degradation defect (Fig. 6C,D). Strikingly, CTD2 mutant protein levels were insensitive to BafA1 treatment, which strongly suggests that SUMOylation-deficient Su(var)2-10 cannot undergo lysosomal degradation.

Finally, we aimed to reveal whether Su(var)2-10 physically interacts with Atg8a *in vivo*, because Atg8a serves as an interaction platform for autophagy cargos and cargo receptors. We indeed detected a specific interaction between certain Su(var)2-10 isoforms and tagged Atg8a by immunoprecipitation from whole fly extracts (Fig. 6E). Collectively, our experiments indicate that interaction of Su(var)2-10 with Atg8a in the nucleus and in the cytoplasm can trigger its autophagic breakdown, which involves Su(var)2-10 SUMOylation.

## Discussion

Here we have uncovered an activation mechanism for the Stat92E transcription factor via autophagy-mediated clearance of a direct repressor. This switch ensures proper glial Stat92E-dependent responses after nerve injury that can contribute to glial reactivity.

The significance of the vital role of autophagy in glia biology is just beginning to unfold^39,40,41^. Neuroprotective microglia in the aging brain show elevated levels of autophagy^42^. Autophagy limits neuroinflammation as it degrades the inflammasome component NLRP3 in microglia to prevent cytokine release and proinflammatory responses^43^. We find that in addition to these functions, autophagy can also positively regulate glial reactivity by impacting a major immune signaling pathway. The autophagic elimination of Su(var)2-10 provides an alternative to canonical JAK-dependent Stat92E regulation that is apparently not employed in glial injury signaling^16^. How injury regulates selective autophagy to remove Su(var)2-10 is an open question, given that bulk autophagy is not elevated in glia after wing injury.

Su(var)2-10 levels increase upon *Atg16* knockdown both in S2 cells and in glia, regardless of injury. The *10xStat92E-GFP* reporter data show that if *Su(var)2-10* is eliminated from glia, Stat92E activity is majorily increased only when flies are simultaneously injured. Su(var)2-10 removal alone is probably insufficient to trigger Stat92E activation because Stat92E activation still depends on tyrosine phosphorylation that is triggered by injury ^16,44^, for its proper function in reactive glia. Therefore, we believe that Stat92E operates as a coincidence detector integrating signals from autophagic removal of Su(var)2-10 and activating phosphorylation for full activation.

The idea of selective autophagic elimination of Su(var)2-10 is also supported by its lysosomal degradation, which does not happen in case of an autoSUMOylation-defective mutant form. SUMOylation regulates many critical developmental and cellular processes and enables the quick responses to environmental and homeostatic changes. Of note, autophagy also contributes to the elimination of SUMOylated Ataxin-3^45^. Ubiquitylation of SUMOylated proteins by SUMO-targeted ubiquitin ligases (STUbLs) can indeed lead to subsequent degradation^46,47^.

Su(var)2-10 colocalizes with Atg8a in both the cytoplasm and nucleus in brain glia. Nuclear Su(var)2-10 also colocalizes with both SUMO and Atg8a. LC3/Atg8a nuclear pools are important for autophagy activation^48,49^. Given that vesicles are excluded from the nucleus, Su(var)2-10-SUMO-Atg8a triple positive structures are likely liquid-liquid phase-separated (LLPS) multimolecular aggregates. SUMOylation regulates stress granule assembly^50^ and other types of phase separation^51^, and the Su(var)2-10 ortholog PIAS1 is predicted to undergo LLPS^52^. The lack of a LIR motif suggests that Su(var)2-10 does not interact directly with Atg8a: it probably requires further labels/interactors that mediate its elimination by autophagy. The Stat92E activation defects in the *Atg8a^LDS^* mutant, the SUMOylation-dependent lysosomal degradation of Su(var)2-10, and its colocalization and biochemical interaction with Atg8a all point to selective autophagic clearance of Su(var)2-10, potentially as part of phase separated particles that are different from p62 bodies.

We find that the alleged antiviral factor *vir-1*^31^ is strongly induced by Stat92E in glia after axon injury, both in the wing nerve and in the brain. *vir-1* expression has been previously documented in glia where it is a target of Gcm, a master regulator of glial cell fate^53^. Importantly, it is also used as a longitudinal glial cell marker in the embryo^54^. A recent study showed strong upregulation of *vir-1* in glia after closed head traumatic brain injury, similarly to antimicrobial peptides^32^. Its expression may be differentially regulated among glial classes since *vir-1* expression defines a distinct subset of fly macrophages^55^. Indeed, *vir-1-GFP* is induced in ensheathing glia and possibly astrocytes in the brain after injury (Fig. 3E) based on the morphology of GFP^+^ cells. Vir-1 features a signal peptide, and it localizes to the extracellular space in the foregut^56^, but its roles are not known.

In the fly brain, *drpr* upregulation after injury is an adaptive response for enhanced debris phagocytosis^57^. We find that *drpr* is highly expressed in wing nerve glia already in uninjured conditions and it is not induced further by wing transection. This could indicate high phagocytic capacity in wing glia, however, axon debris clearance takes substantially longer in the wing nerve than in antennal lobe glomeruli of the brain^15,58^. Thus, Drpr levels may not readily predict actual phagocytic activity in glia.

Glial reactivity in *Drosophila* likely relies to a great extent on Stat92E signaling, because in the CNS, Stat92E induces *drpr* and *matrix metalloproteinase 1* (Mmp1) that promote phagocytosis and access to cell debris via ECM degradation^16,17^. Furthermore, *stat92e*-deficient glia do not even extend processes towards injured axons^16,17^. Correspondingly, phosphorylated STAT3 is a marker for reactive astrocytes in mice^59^. Illustrating the complex functionality of this pathway in mammals, STAT3-dependent transcription has a protective role during traumatic brain injury, motor neuron injury, spinal cord injury and neonatal white matter injury, while it contributes to degeneration in AD models^59^. Accordingly, glial Stat92E activation is also observed in an amyloid-beta expressing *Drosophila* model^60^. We find that selective autophagy promotes Stat92E signaling and thereby it could assist functional recovery after injury. In line with this, autophagy deficiency in microglia predisposes to increased neurodegeneration and demyelination in a multiple sclerosis (MS) model^42^. Based on our findings, we expect that the benefits of autophagy stimulation in the nervous system will partly manifest through STAT transcription factor-driven glial reactivity.

## Acknowledgements

Funding

This research was supported by:

- the Momentum/Lendület Grant of Hungarian Academy of Sciences (LP2023-6) (GJ)
- the Young Researchers’ Excellence Programme of the National Research, Development and Innovation Office (NRDIO) (FK132183) (AS)
- the Janos Bolyai Research Scholarship of the Hungarian Academy of Sciences (BO/00078/18) (AS)
- the NRDIO New National Excellence Programme (ÚNKP-20-5, ÚNKP-19-4, ÚNKP-18-4) (AS)
- the NRDIO grant KKP129797 (GJ)
- the NRDIO grant PD145868 (AJ)
- the NRDIO grant GINOP-2.3.2-15-2016-00032 (GJ)
- and the Biotechnology National Laboratory program of the National Research, Development and Innovation Office (NKFIH-871-3/2020) (GJ).
- the National Talent Programme - Scholarship for the Young Talents of the Nation (NTP-NFTÖ-22-B-0055) (VV),
- the New National Excellence Programme (ÚNKP-23-3 -SZTE-537) (VV)
- the University Research Fellowship Programme (EKÖP-24-3-SZTE-602) (VV)
- and the National Academy of Scientist Education Program of the National Biomedical Foundation under the sponsorship of the Hungarian Ministry of Culture and Innovation (ZE, AS and GJ).

We thank László Tirián and Julius Brennecke for the mouse anti-Su(var)2-10 antibody, Giacomo Cavalli for the rabbit anti-SUMO antibody, Marc Freeman for *dee7-Gal4* and other stocks, Jean-Luc Imler for *vir-1-GFP,* Ioannis Nezis for the *Atg8a^LDS^* stock, Melinda Bence and Miklós Erdélyi for *Su(var)2-10* RNAi stocks, the Su(var)2-10 expression constructs and further support, Zoltán Hegedűs from the Laboratory of Bioinformatics for bioinformatic analyses, the Bloomington Stock Center and the Vienna *Drosophila* Resource Center for fly stocks, the Developmental Studies Hybridoma Bank for antibodies, Szilvia Bozsó and Ildikó Erdődi for technical assistance. We also acknowledge the provided microscopy support of the Cellular Imaging Laboratory (HUN-REN Biological Research Centre). We are grateful to Gábor Csordás, Arindam Bhattacherjee, András Blastyák and Tamás Maruzs for fruitful discussions. Fig. 3A, 7, S1A and S2A were created by BioRender.com.

## Author contributions

Conceptualization: V.V., A.S. and G.J.; Investigation: A.S., V.V., Z.E., E.F.J., A.S.C., A.J., M.C., D.F.B., A.R.G. and G.J. Resources: A.J., Z.E., D.F.B., A. R. G.; Results interpretation: A.S., V.V., Z.E., E.F.J., A.S.C., A.J., M.C., D.F.B., A.R.G. and G.J. Writing: V.V., Z.E., A.S. and G.J.; Manuscript revision: V.V., Z.E., A.S., A.S.C., and G.J.; Supervision: A.S. and G.J.; Funding acquisition: A.S. and G.J.

## Data availability

All data needed to evaluate the conclusions in this paper are present in the paper and its Supplementary Information. Other data associated with the article such as raw data are available upon request.

## Materials availability

All materials, *Drosophila* stocks and related information are available from the corresponding authors upon reasonable request.

## Ethics statement

The research presented here uses the species *Drosophila melanogaster* for which no ethical approval is required in the HUN-REN Biological Research Center, Institute of Genetics or by Hungarian authorities. Maintenance of transgenic *Drosophila* melanogaster at the Institute is regulated by license No. BGMF/867-9/2022 of the Gene Technology Authority Registry, Ministry of Agriculture of the Hungarian Government.

## Materials and methods

### *Drosophila* stocks and maintenance

Flies were kept at 25°C on a medium containing cornmeal, yeast, agar and dextrose with Nipagin as a preservative in reusable glass vials and bottles. For our experiments, we injured male *Drosophila melanogaster* that were 2–7 days old after eclosion. During our experiments, we favored male flies to avoid technical inconvenience related to egg laying and to prevent larvae from making the medium too liquid and sticky. The animals were anesthetized by carbon dioxide. Progeny of *repo-GS* crossed with RNAi lines were raised on normal food during development, then males were kept for 5-7 days on food supplemented with 25 μg/ml RU486/mifepristone (Acros Organics) before axotomy and thereafter during the course of the experiment. Injury was carried out on 7-12 day old flies. The supplemented medium used for experiments using *vir-1-GFP* contained 125 μg/ml RU486.

The following stocks were ordered from the Bloomington *Drosophila* Center: *repo-Gal4 (7415), w^1118^*, *daughterless-Gal4 (95282)*, *vasa-Cas9* (*51323*), *TRE-EGFP (59010), firefly luciferase^JF01355^, Atg8a^TKO.GS04635^*(*80828*), *Su(var)2-10^JF03384^ (29448), Su(var)2-10^HMS00750^ (32956), UASp-mCherry::Atg8a (37750), UASp-GFP-mCherry-Atg8a (37749), UAS-TransTimer (93411), drpr^MI07659^ (843909), UAS-2xEYFP (60291), vas-int, w[*], hs-Cre (60299), pP{lox(Trojan-GAL4)x3} (60310)*^61^*, Su(var)2-10^MI03442^ (37190), stat92E^GL00437^ (35600)*, *10XStat92E-ΕGFP (26197), UAS-NLS::GFP (4775)* and *10XUAS-IVS-myr::tdTomato (32221)*.

The following stocks were provided by the Vienna *Drosophila* Resource Center: *Rubicon^KK108247^, Atg13^KK100340^, Atg16^KK10232^* and *ref(2)P^KK105338^*. The *Atg5^5cc5^* ^62^*, Atg8a^LDS^* ^27^, *Atg5^5cc5^; Atg5::3xHA* ^24^ stocks were formerly created and published. *Atg101^Δ6h^* was contributed by Wanzhong Ge (Institute of Genetics, Zhejiang University School of Medicine, Hangzhou, China)^63^. The following stocks received as gifts were much appreciated: *repo-GeneSwitch* was kindly shared by Véronique Monnier (Université de Paris, BFA Unit of Functional and Adaptative Biology, UMR 8251, France). *dee7-Gal4* (*drpr* enhancer-*Gal4*) was kindly contributed by Marc Freeman (Vollum Institute, Oregon Health & Science University, USA), the *Atg8a^LDS^* stock by Ioannis Nezis (University of Warwick, UK) and *vir-1-GFP.2.6 (vir-1-GFP)* by Jean-Luc Imler (IBMC, University of Strasbourg, France). Knockdown efficiency of Atg gene RNAi lines were validated previously^26^.

The *Atg8a^null^* frameshift mutant was generated by CRISPR-Cas9 editing. To generate this null allele (*Atg8a^Δ12^*), we used the *Atg8a^TKO.GS04635^* gRNA line and crossed it with *vasa>Cas9* transgenic flies. We screened candidate mutant lines by PCR and Sanger sequencing and identified the *Atg8a^Δ12^* allele that carries an 8 bp deletion based on sequencing data (Fig. S2A), which deletion causes frameshift in all isoforms of the Atg8a gene.

The 2xmApple::Atg8a construct was cloned into the pACU vector (Addgene #58373). mApple sequences were amplified from the pCDH-TMEM106b-mApple vector (Addgene #179385). The PCR products which were used for the construct assembly with the NEBuilder® HiFi DNA Assembly Master Mix (New England Biolabs) are listed in the Supplementary Table 2. We validated the constructs by Sanger sequencing and we were able to detect mApple fluorescence in induced transfected S2 cells cotransfected with pMT-Gal4. The transgenic *Drosophila* stock was established by the *Drosophila* Injection Service in the Biological Research Center, Szeged (Hungary). The transgene was inserted into the *attP40* (2L) landing site.

A MiMIC transgene (*MI07659*) located in intron 6 of *drpr* was converted into a *drpr-Gal4* fusion gene using a *Gal4* donor, so that Gal4 is produced instead of most of the *drpr* coding sequence. We followed the protocol described in Diao et al. 2015^61^ by crossing in recombinase and donor transgenes. A donor cassette containing a splice acceptor (*SA*), the *Gal4* to be encoded in the correct frame and finally a stop codon and polyadenylation site between two *attB* sites was mobilized and circularized by heat shock-inducible Cre. Integration into *MI07659* was achieved by vas-dΦC31. Candidate progeny were screened for *drpr* expression by crossing to *UAS-2xEYFP*. Integration into *drpr* was validated by PCR. *drpr-Gal4* is expressed in glia in the CNS and PNS in addition to other tissues.

### Rearing axenic stocks for RT-qPCR

We placed approximately 100 animals from the used strains into a cylinder containing black egg-laying agar coated with yeast for one night. Embryos were dechorionated in 4-5% hypochlorite on a filter for 1 minute followed by washing with flowing distilled water. Finally, we removed the filter fabric, placed it on a paper towel, and transferred the eggs into new vials containing a nutrient medium mixed with antibiotics already described^64^. Autoclaved cornmeal-yeast-agar-dextrose–Nipagin medium was mixed with 0.25 mg/ml ampicillin, tetracycline, streptomycin, and 1 mg/ml kanamycin^64^. Flies were kept for at least 3 generations on antibiotics. The concentration of the antibiotics were reduced to half 20 days before sample collection to increase the amount of progeny. The food cocktail preparation and fly transfer took place in a laminar box using only autoclaved and sterile tools.

### Wing injury

In all experiments we carried out in *Drosophila,* male flies were used. In the wing injury assay, we used the axotomy model developed by Fang et al.^23^ and Neukomm et al^22^. In accordance with the model (S1A), we removed half of the wing of flies from the distal end with a straight cut across using microsurgical scissors (Fine Science Tools). For each individual, we only cut one wing, leaving the other intact as an uninjured control. The injured animals were then kept in vials at 25 °C. Three days after the injury (3 dpi), a strong *10xStat92E-GFP* signal was already observable (Fig. S1D,E).

### Antennal ablation

The antennae were ablated 1 day prior to brain dissection by pulling on with a forceps^15^, and the injured flies were kept in vials at 25°C.

### Immunostaining of adult brains

Immunostaining of adult brains was performed as described with some modifications^65^. Adult brains were dissected in ice-cold phosphate buffered saline (PBS) and placed immediately in 4% paraformaldehyde in PBS with 0.3% Triton X-100 (PBT) on ice. Brains were fixed for 1 h at 25 °C. After two quick rinses with PBT, brains were washed three times 20 min each. Following blocking in 5% fetal bovine serum (FBS) in PBT for 1 h at RT (RT), brains were incubated with the primary antibody (mouse anti-Su(var)2-10^66^ 1:100, rabbit anti-GFP 1:1000 Thermo Fisher Scientific A-11122, rabbit anti-SUMO^67^ 1:200, rabbit anti-mCherry NBP2-25157, Novis Biologicals 1:500) for 3 days at 4 °C in 5% FBS in PBT. The SUMO antibody was gently provided by Giacomo Cavalli (Institute of Human Genetics (IGH), Montpellier, France) and the Su(var)2-10 antibody by Julius Brennecke (IMBA, Vienna, Austria). Washes were done as before and brains were incubated with the fluorescently labelled secondary antibody (goat anti-rabbit Alexa Fluor 488, A-11034, goat anti-mouse Alexa Fluor 568, A-11031, from Thermo Fisher Scientific, all 1:1000 diluted) in 5% FBS in PBT 2 days at 4 °C in darkness. The secondary antibody was removed by two quick rinses, and DAPI (1:15000) was put into the following first wash with PBT. Brains were washed three times 20 min each. After the washes, the brains were mounted in Vectashield (Vector Laboratories, H-1000-10), and the samples were kept in darkness at 4°C until imaging.

### Imaging

For imaging reporter signals in the wing, we focused on a specific region inside the L1 vein proximal to the injury site, a curvature from the wing margin towards the inner area of the wing, containing axons and glial cells (Fig. S1A). We analyzed the samples at room temperature (RT) immediately after dissection and mounting into Halocarbon Oil 27 (Sigma-Aldrich, H8773), with an Axio Imager.M2 structured illumination microscope (Zeiss) and the ORCAFlash4.0LT CMOS camera (Hamamatsu) using a Zeiss PlanApochromat 63x/1.40 NA objective. Optical slices were created with a Zeiss ApoTome.2 device. The illumination was provided by the CoolLED pE-4000 system. We captured the images using the Zeiss ZEN program.

For measuring the Su(var)2-10 signal in Su(var)2-10 and NLS-GFP co-immunostained brains, the Visitron VisiScope Spinning Disk Confocal Microscope (Visitron Systems GmbH, Puchheim, Germany) was used. The optical slices were created with an Olympus LUCPlanFl 40x NA 0.60 objective for quantification, while magnified images shown were captured with an Olympus PlanApo 60x NA 1.42 (oil) objective. Detection was based on an Andor Zyla 4.2 PLUS camera.

For the three channel fluorescence images, 405 nm excitation with 405/488/561 tri band emission filter, 488 nm excitation with 525/50 emission filter, and 561 nm excitation with 605/70 emission filter were used with a triple band 405/488/561 dichroic mirror. For the four channel fluorescence images, 405 nm excitation with 405/488/561 tri band emission filter, 488 nm excitation with 525/50 emission filter, 561 nm excitation with 605/70 emission filter, and 640 nm excitation with 700/75 emission filter were used with a quad band 405/488/561/640 dichroic mirror. Channels were acquired in sequential mode, after the performed Z sectioning. We used a disk with 50 μm pinholes. For *vir-1-GFP* signal detection in brain, we used an LSM800 (Zeiss) inverted laser scanning confocal microscope. The brains were imaged at RT with a Zeiss Plan-Apochromat 40x/1.3 NA oil immersion objective with a Z-step of 2.00 μm. The same microscope settings (illumination, exposure time, laser intensity) were used for all conditions and genotypes throughout one experiment.

### RNA purification and RT-qPCR

For RNA isolation, we collected around 150-200 wings from axenic flies per sample using only autoclaved tools with autoclaved wiping papers placed on the fly sorting pad previously washed with 70% ethanol. The wings were swept in TRI Reagent (Zymo Research)-containing microcentrifuge tubes, using a funnel. We homogenized the samples directly after putting them into TRI Reagent. Total RNA was purified with the Direct-zol RNA Microprep (Zymo Research). DNAse I digestion was also applied. 76 ng total RNA was reverse transcribed in 20 μl reaction volume with the RevertAid First Strand cDNA Synthesis Kit (Thermo Scientific). qRT-PCR was performed in 20 μl reactions in technical triplicates using the PerfeCTa SYBR Green FastMix (Quantabio) with 0.5 μl cDNA and cycled on a Rotor-Gene Q qPCR machine (Qiagen) running Rotor-Gene software 2.3.1.49 (Qiagen), with the following program: 95 °C, 3 min; 45 cycles of 95 °C, 20 s, 60 °C, 20 s and 72 °C, 20 s followed by melting curve analysis. The ΔΔCt method was employed to normalize the data, with *Ribosomal protein L32* (*RpL32*, also known as *rp49*) serving as the internal control. All primers were designed with Primer-BLAST (https://www.ncbi.nlm.nih.gov/tools/primer-blast) with amplicon length set to 70– 150 bp. The used melting temperature was 60 °C. One primer spanned an exon-exon junction in all cases. Primers used for RT-qPCR are listed in the Supplementary Table 2. We normalized the qPCR datasets to RpL32. The ΔCt-derived expression values were adjusted by a common scaling factor to ensure that the average of control values equalled 1 or 100.

### *Drosophila* S2R+ cell culture

*Drosophila* S2R+ cells were cultured in 1x Schneider’s *Drosophila* media (GIBCO) supplemented with 10 % fetal bovine serum (GIBCO), penicillin-streptomycin solution (HyClone) in 25-cm^2^ T-flasks (Thermo Fisher Scientific) at 27 °C. The cells (∼8×10^5^) were seeded in a six well-culture plate and 24 h later were transiently co-transfected with *pUASp-Su(var)2-10-WT::GFP*^38^*, pUASp-Su(var)2-10-CTD2mut::GFP* and *pMT-Gal4* using the TransIT-Insect (MIR 6100, Mirus Bio) transfection reagent based on the manufacturer’s recommendations. S2R+ cells were transfected with 2500 ng of each plasmid. Twenty-four hours after transfection, 0.5 mM of CuSO_4_ was added to the cells to induce Gal4 expression from the metallothionein promoter. 24h after induction, cells were treated with 100 nM Bafilomycin A1 (B1793, Sigma), and incubated for 4 h. Cell pellets were stored at −80 °C.

### Immunoprecipitation

Extracts were prepared from 0.5 ml of 3-7 days old whole flies from *da-Gal4>UAS-myr-tdTomato* and *da-Gal4>UAS-mCherry::Atg8a* genotypes. Flies were homogenized in 0.5 ml extraction buffer (50 mM Tris pH = 7.4, 150 mM NaCl, 2 mM EDTA, 0.5% Nonidet P-40, 5% glycerol, Pierce Protease Inhibitor Tablet (A32963, Thermo Fisher Scientific) and Halt Protease and Phosphatase Inhibitor Cocktail (78442, Thermo Fisher Scientific) on ice in a Potter homogenizer. Following centrifugation (15000 g, 15 min, 4 °C), the protein concentration in the supernatant was measured with Pierce Bradford Plus reagent (23238, Thermo Fisher Scientific). 5 mg extract was precleared by rotating with 50 μl of Dynabeads M-280 Streptavidin (11206D, Thermo Fisher Scientific) for 1h at 4 °C to absorb bead-binding nonspecific proteins and immunoprecipitated using 50 μl of RFP-Trap magnetic Agarose (rtma, ChromoTek) for another 1h at 4 °C. Subsequently, beads were washed with extraction buffer 3 times for 10 min each. Beads were eluted in 20 μl of 2x Laemmli sample buffer without DTT for 5 min at 100 °C and diluted and supplemented with DTT. Half of the eluted sample was loaded on gels.

### Western blotting

S2 cell pellets were lysed on ice for 30 min in three times packed volume RIPA buffer (50 mM Tris pH = 8, 150 mM NaCl, 1% Nonidet P-40, 0.5% sodium deoxycholate, 0.1% SDS, Pierce Protease Inhibitor Tablet (A32963, Thermo Fisher Scientific), Halt Protease and Phosphatase Inhibitor Cocktail ((78442, Thermo Fisher Scientific)) and cleared by centrifugation at 4 C. Protein concentration in the supernatant was measured with Pierce Bradford Plus reagent (23238, Thermo Fisher Scientific). 5 μg total protein was loaded onto 8% polyacrylamide gels. L3 stage homozygous larvae from the *Su(var)2-10^MI03442^*/*CyO-GFP* and *w^1118^* genotypes were collected from black agar plates after synchronized egg laying and homogenized in 6 μl Laemmli buffer per μl larva with a pipette tip, then with a motor pestle for 20 sec. 20 μl extract was loaded on 8% polyacrylamide gels. After overnight blotting on Immobilon^®^-FL PVDF Transfer Membrane (Merck), membranes were blocked with Intercept (TBS) Blocking Buffer (LI-COR) for 1 h at RT. The following primary antibodies were used: rabbit anti-GFP (Thermo Fisher Scientific A-11122), rabbit anti-mCherry (NBP2-25157, Novis Biologicals), mouse anti-Su(var)2-10^66^, mouse anti-α-tubulin (Developmental Studies Hybridoma Bank AA4.3) and rabbit anti-β-actin (Thermo Fisher Scientific PA5-85271) all 1:1000, diluted in Blocking Buffer and incubated for 1 h at RT. Secondary antibodies were the following: anti-mouse IRDye 800CW (926-32210, LI-COR) and anti-rabbit IRDye 680RD (926-68071, LI-COR) 1:15,000, diluted in Blocking Buffer supplemented with 0.02% SDS and 0.2% Tween-20 and incubated for 1h at RT. Dried blots were imaged and quantified in an Odyssey Clx instrument (LI-COR).

### Image analysis

The signal intensity coming from the fluorescent images were quantified with Fiji (ImageJ 2.9.0, v1.54h). For assessing *TRE-EGFP*, *10xStat92E-GFP* and *vir-1-GFP* signal intensity levels in wings, we measured a 174 x 54 pixel area in each case. When quantifying the intensity of the vir-1-GFP signal in the brain, intensity data were generated from both optic lobes within a 225 x 37 pixel area. To determine the level of Su(var)2-10 within the nuclei, we randomly selected 10 glial cells surrounding the AL marked by *repo-Gal4 > NLS-GFP* from each brain (except in cases where fewer cells were found). The nuclei were outlined using freehand selection. To measure *drpr* activity in the wing by using *drpr-Gal4 > UAS-TransTimer* flies, we selected an area of 800 x 200 pixel and measured all the nuclei intensities inside the rectangles. The nuclei were outlined by freehand selection. In *dee7-Gal4 > UAS-TransTimer* brains, we used a 1300 x 1300 pixel area to count red puncta. In *dee7-Gal4 > UAS-TransTimer* wings, a 600 x 140 pixel sized rectangular area was selected for counting red puncta. All fluorescent signals detected in the area of interest were normalized to the background with identical area sizes. Western blot bands were quantified by the Image Studio 5.2 software (LI-COR). Su(var)2-10 signals were normalized to α-tubulin levels.

### Statistical analysis

Experiments were independently repeated at least two times with differing biological samples, with similar results. No data points were excluded. The quantified results from image analysis are represented by truncated violin plots with median and quartiles containing all data points. RNA and protein measurements are presented as mean and standard deviation. The statistical analyses were performed with GraphPad Prism 8.0.1 and 10. We first checked the normal distribution within each data group by running the Shapiro-Wilk normality test (α = 0.05). When conducting pairwise comparisons on normally distributed data, we performed unpaired, two-tailed Student’s *t*-test, while for non-normally distributed data, the unpaired, two-tailed Mann-Whitney test was applied. For the comparison of multiple datasets of normal distribution, one-way ANOVA with Holm- Šídák’s test for multiple comparison correction was used as in Fig. 6D. No power calculations were performed. For the experiments, we always used a sample size that gave reproducible results, similarly to a relevant article^11^. Western blot and qPCR results were evaluated with n = 3 or 4 biological replicates.

