## Supplemental Material for "Selective autophagy fine-tunes Stat92E activity by degrading Su(var)2-10/PIAS during glial injury signaling in *Drosophila*"

#### Fig. S1

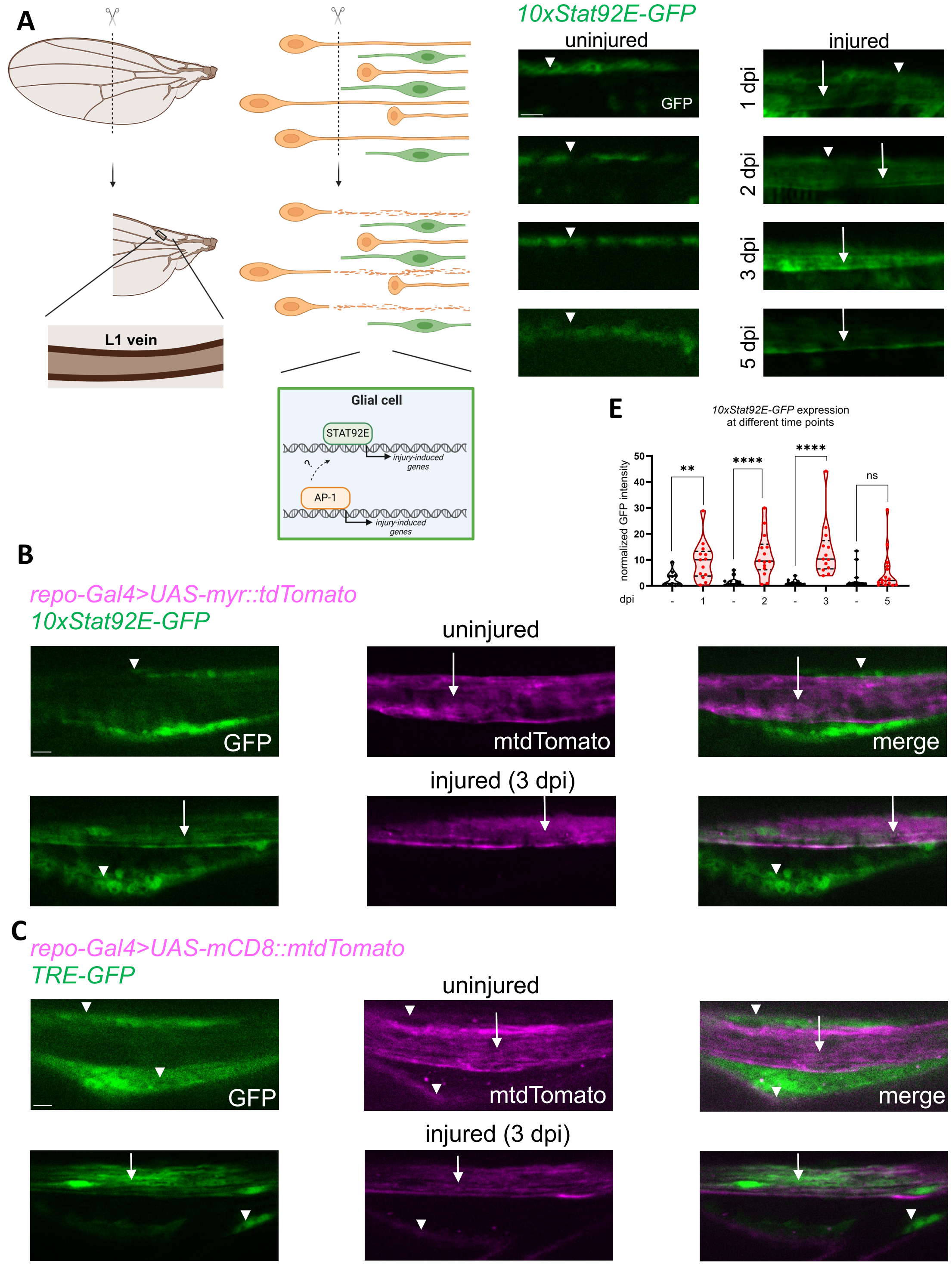
A
L1 vein
Glial cell
STAT92E
AP-1
injury-induced genes
injury-induced genes
B
repo-Gal4>UAS-myr::tdTomato
10xStat92E-GFP
uninjured
GFP
mtdTomato
merge
injured (3 dpi)
C
repo-Gal4>UAS-mCD8::mtdTomato
TRE-GFP
uninjured
GFP
mtdTomato
merge
injured (3 dpi)
D
10xStat92E-GFP
uninjured
injured
GFP
1 dpi
2 dpi
3 dpi
5 dpi
E
10xStat92E-GFP expression
at different time points
normalized GFP intensity
dpi
**
****
****
ns
0
10
20
30
40
50
-
1
-
2
-
3
-
5

#### Fig. S2

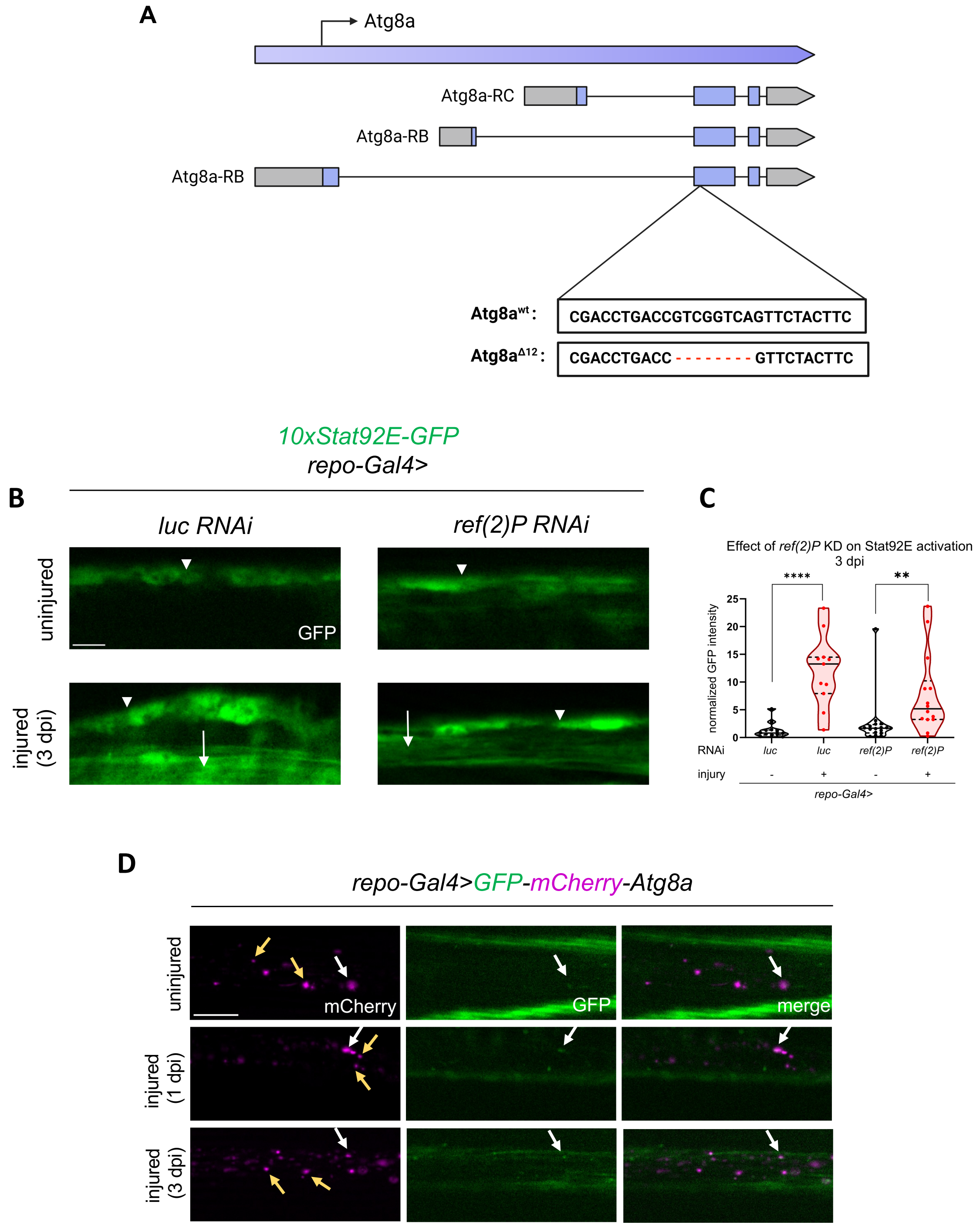

A
Atg8a
Atg8a-RC
Atg8a-RB
Atg8a-RB
Atg8awt :
CGACCTGACCGTCGGTCAGTTCTACTTC
Atg8aΔ12 :
CGACCTGACC - - - - - - - - GTTCTACTTC
10xStat92E-GFP
repo-Gal4>
B
luc RNAi
ref(2)P RNAi
uninjured
injured (3 dpi)
GFP
C
Effect of ref(2)P KD on Stat92E activation
3 dpi
****
**
normalized GFP intensity
25
20
15
10
5
0
RNAi
luc
luc
ref(2)P
ref(2)P
injury
-
+
-
+
repo-Gal4>
D
repo-Gal4>GFP-mCherry-Atg8a
uninjured
injured (1 dpi)
injured (3 dpi)
mCherry
GFP
merge

#### Fig. S3

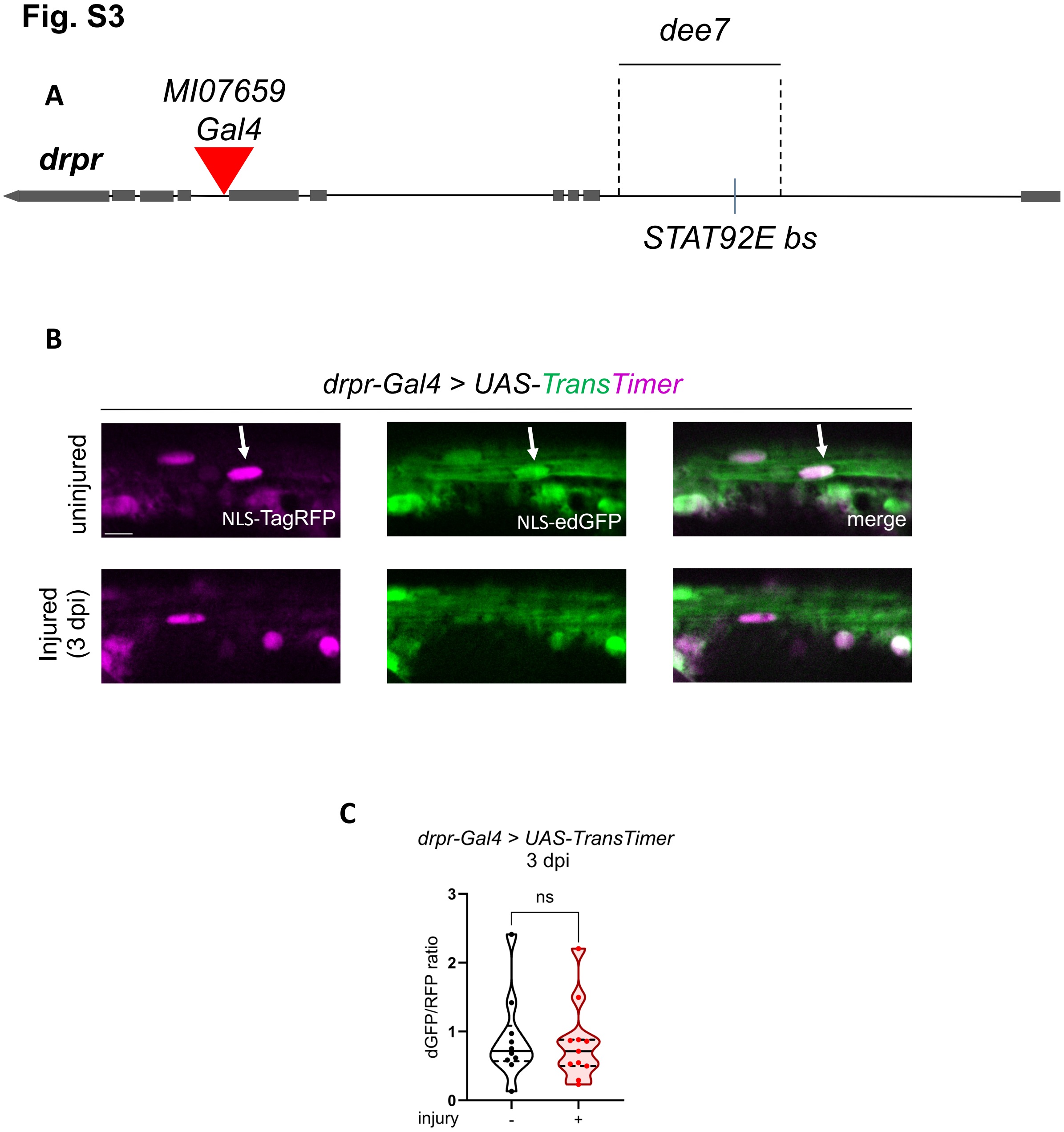
A
dee7
MI07659
Gal4
drpr
STAT92E bs
B
drpr-Gal4 > UAS-TransTimer
uninjured
Injured
(3 dpi)
NLS-TagRFP
NLS-edGFP
merge
C
drpr-Gal4 > UAS-TransTimer
3 dpi
ns
dGFP/RFP ratio
0
1
2
3
injury
-
+

Fig. S4

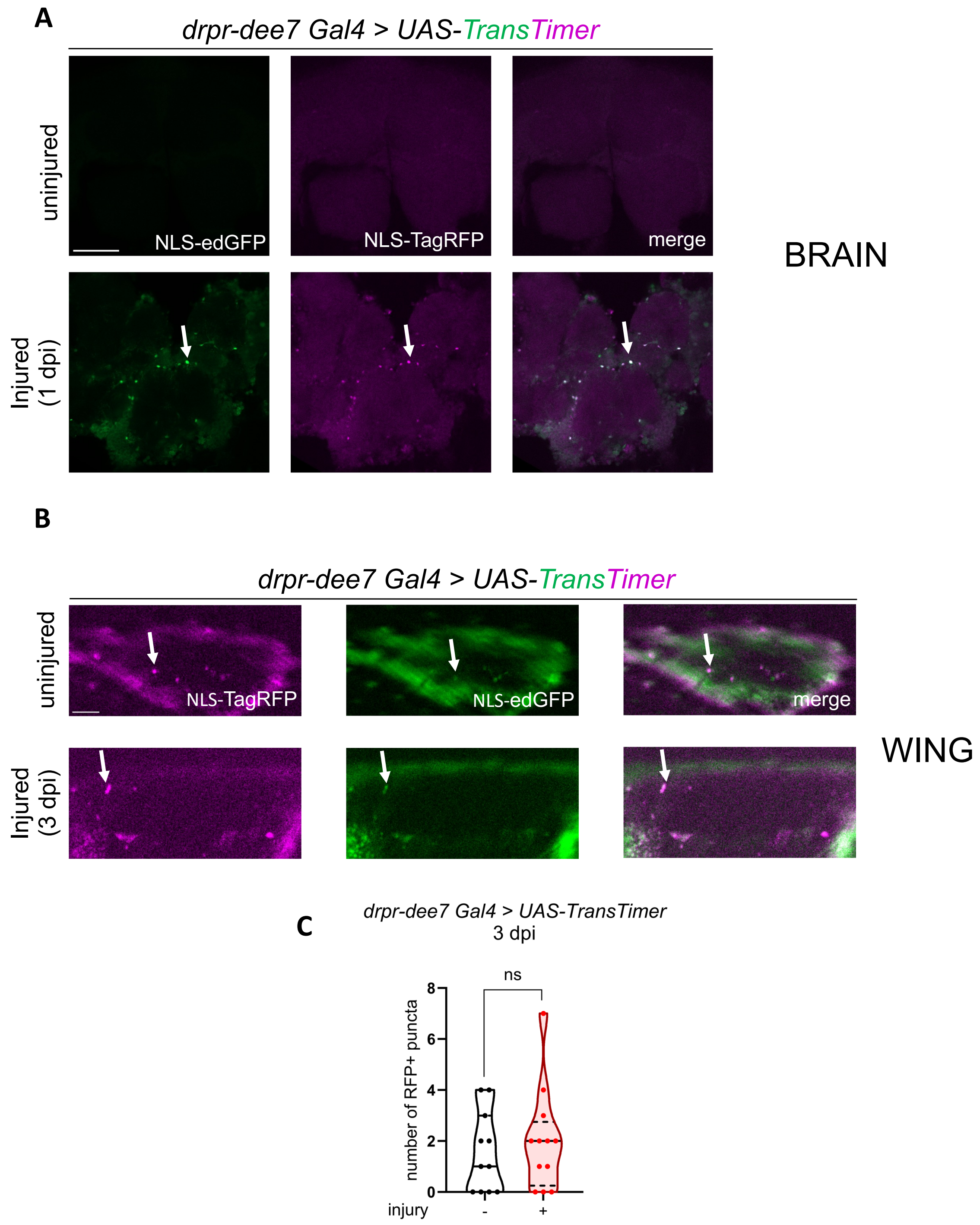
A
drpr-dee7 Gal4 > UAS-TransTimer
uninjured
Injured (1 dpi)
NLS-edGFP
NLS-TagRFP
merge
BRAIN
B
drpr-dee7 Gal4 > UAS-TransTimer
uninjured
Injured (3 dpi)
NLS-TagRFP
NLS-edGFP
merge
WING
C
drpr-dee7 Gal4 > UAS-TransTimer
3 dpi
ns
number of RFP+ puncta
0
2
4
6
8
injury
-
+

#### Fig. S5

**A**

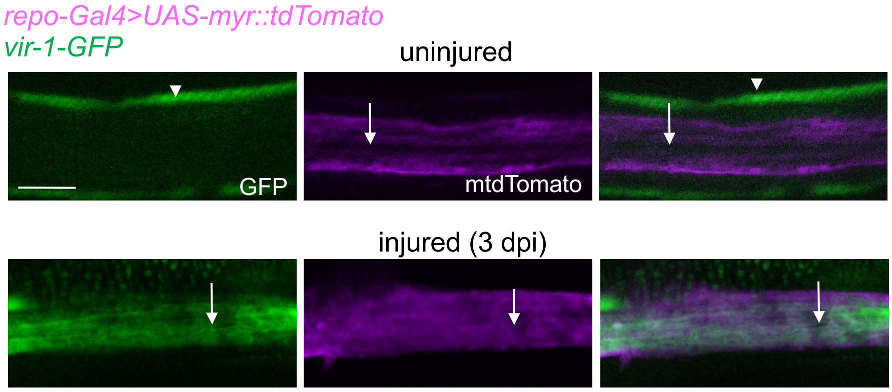

repo-Gal4>UAS-myr::tdTomato
vir-1-GFP
uninjured
GFP
mtdTomato
injured (3 dpi)

**B**

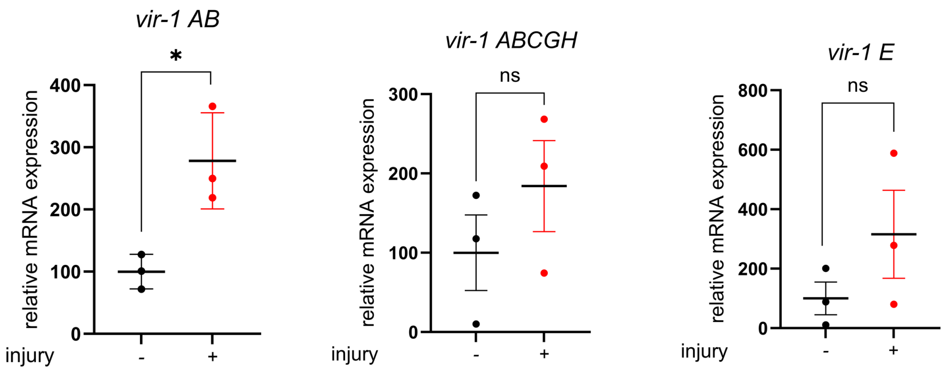

vir-1 AB
*
relative mRNA expression
400
300
200
100
0
injury
-
+
vir-1 ABCGH
ns
relative mRNA expression
300
200
100
0
injury
-
+
vir-1 E
ns
relative mRNA expression
800
600
400
200
0
injury
-
+

#### Fig. S6

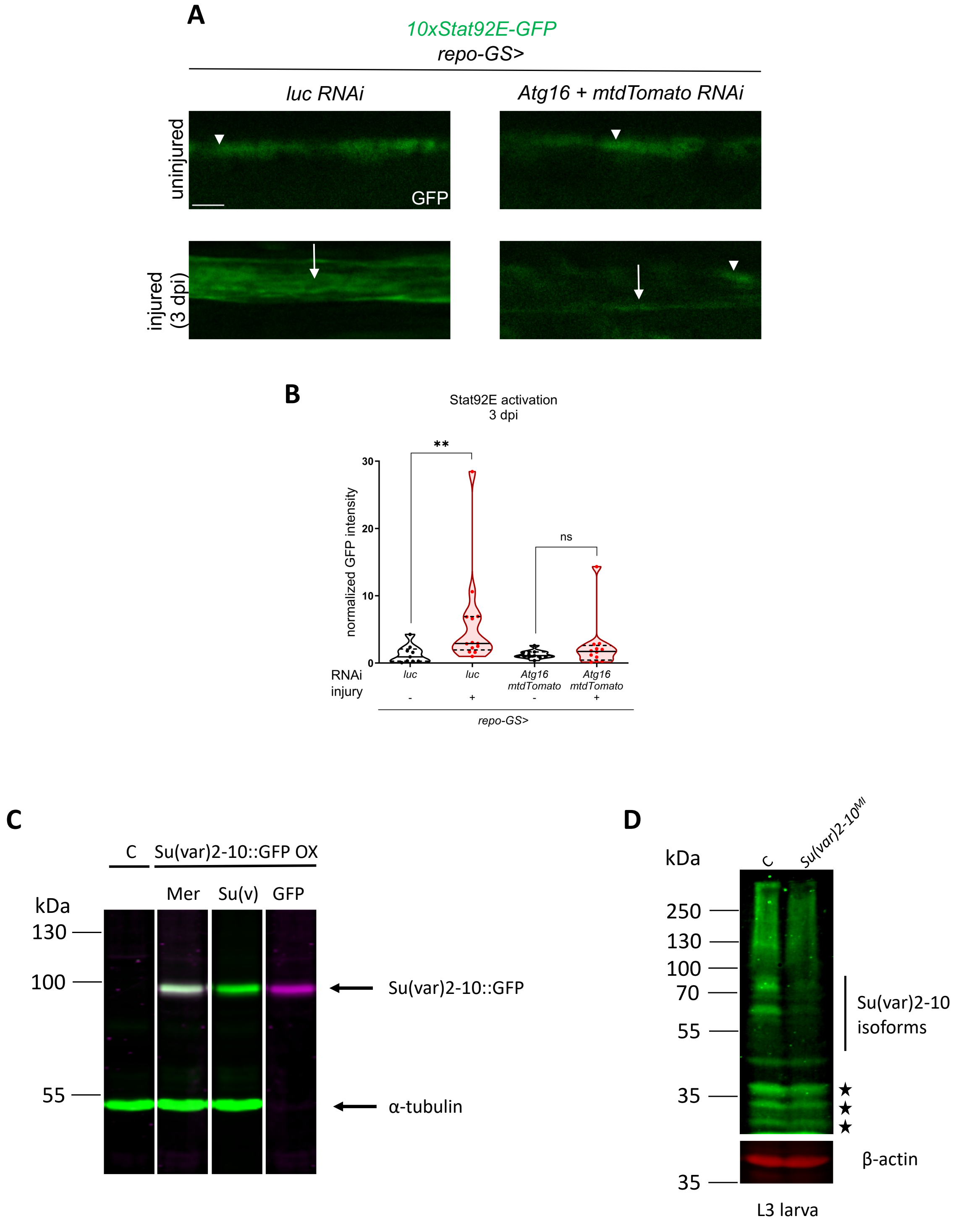
A
10xStat92E-GFP
repo-GS>
luc RNAi
Atg16 + mtdTomato RNAi
uninjured
injured (3 dpi)
GFP
B
Stat92E activation
3 dpi
normalized GFP intensity
30
20
10
0
**
ns
RNAi
injury
luc
luc
Atg16 mtdTomato
Atg16 mtdTomato
-
+
-
+
repo-GS>
C
C
Su(var)2-10::GFP OX
Mer
Su(v)
GFP
kDa
130
100
55
Su(var)2-10::GFP
α-tubulin
D
C
Su(var)2-10MI
kDa
250
130
100
70
55
35
35
Su(var)2-10 isoforms
β-actin
L3 larva

### Supplementary Figure Legends

**Figure S1. Stat92E and AP-1 transcriptional reporters are activated in wing glia after injury**

(A) Schematic representation of the injury-induced glial reactivity in *Drosophila* wing. After transecting the wing, severed axons (yellow) will undergo Wallerian degeneration and trigger reactivity in circumambient glia (green). This results in transcriptional activation by signalling pathways. It is known that transcription factors AP-1 of the JNK pathway and Stat92E are activated after axonal injury in CNS glia. The rectangular shaped outline indicates the imaged area within the L1 vein nerve. (B,C) Fluorescent microscopic images of uninjured and injured wing nerves 3 dpi. Single optical slices are shown in each case. *repo-Gal4* driven myr::mtdTomato colocalizes with the signal from the *10xStat92E-GFP* and the *TRE-EGFP* reporters in glia. (D) Single-slice images of wings with *10xStat92E-GFP* signal in glia 1, 2, 3 and 5 dpi show the activity of the Stat92E transcription factor. (E) Quantitative analysis of *10xStat92E-GFP* signal intensity at 4 different time points shown in (D). Truncated violin plots are shown with median and quartiles. Unpaired, two-tailed Mann–Whitney test was used for statistics. **** $p < 0.0001$, ** $p = 0.0024$, ns = 0.1627. n = 12, 13, 13, 14, 13, 13, 15, 16. The arrows point to glial reporter expression, while the arrowheads denote the epithelial expression surrounding the veins. Scale bar: 5 µm.

**Figure S2. The Ref(2)P/p62 cargo receptor is not involved in Stat92E activation and the volume of bulk autophagy is not increased in glia after wing injury**

(A) Schematic illustration shows the site and sequence of the mutation in the CRISPR-Cas9 generated *Atg8a*$^{\Delta12}$ flies which causes a frameshift in all transcript variants. (B) Single optical slices of wing nerves with glial *ref(2)P* silencing in a *10xStat92E-GFP* background. RNAi expression was induced by *repo-Gal4.* The arrows show the reporters expressed in glia, while the arrowheads denote the epithelium surrounding the veins. Scale bar: 5 µm. (C) Quantitative analysis of *10xStat92E-GFP* signal in *ref(2)P* RNAi. Truncated violin plots are shown with median and quartiles. The analyses were carried out by unpaired, two-tailed Mann-Whitney test. **** $p < 0.0001$. ** $p = 0.0013$. n = 12, 11, 16, 14. (D) Wings of *repo-Gal4>UAS-GFP-mCherry-Atg8a* flies were transected and imaged at the indicated time points. White arrows point to double GFP+ mCherry+ vesicles while yellow arrows point to only mCherry+ vesicles. mCherry is stably fluorescent in lysosomes while GFP is rapidly quenched, therefore autolysosomes are exclusively mCherry+ while other autophagic structures are doubly labelled. Scale bar: 10 µm.

**Figure S3. *drpr* is not induced by injury in wing glia**

(A) Schematic representation of the *drpr* locus. *dee7* bears a Stat92E-responsive enhancer including a functional Stat92E binding site (bs). *MI07659* Trojan Gal4 conversion reflects endogenous *drpr* expression. (B) *drpr-Gal4* drives TransTimer (NLS-TagRFP and NLS-ed(enhanced destabilized)GFP separated by a 2A sequence) in the wing nerve. At 3 dpi, no major change is observed in the number of double GFP+RFP+ nuclei that reflects *drpr* promoter-enhancer activity. The arrows point to the nuclei. Scale bar: 5 µm. (C) Truncated violin plots with median and quartiles are shown for the quantification of the NLS-edGFP / NLS-TagRFP ratio in nuclei in *drpr-Gal4* driven *UAS-TransTimer* flies without injury and 3 dpi. The analysis was carried out by unpaired, two-tailed Mann-Whitney test. ns = 0.8094. n = 10, 11.

**Figure S4. *drpr* enhancer *dee7* is not induced by injury in wing glia**

(A) *dee7-Gal4* drives *UAS-TransTimer* in the brain. The central brain is shown 1 day after antennal ablation. In accordance with the published results[16], *dee7* enhancer activity robustly increases after injury reflected by the appearance of double GFP+RFP+ nuclei. The arrows denote the RFP+ GFP+ puncta. Scale bar: 50 µm. (B) *dee7-Gal4* drives *UAS-TransTimer* in the wing nerve in uninjured and injured condition. As opposed to the brain, there is a basal activity of *dee7* in the wing nerve that does not appear to increase after wing transection. The arrows point to the RFP+ and / or GFP+ puncta. Scale bar: 5 µm. (C) Statistical analysis of the number of RFP+ puncta in uninjured wings and 3 dpi, shown in (B). Truncated violin plots with median and quartiles are shown. The statistics were carried out by unpaired, two-tailed Mann-Whitney test. ns = 0.6575. n = 11, 12.

**Figure S5. *vir-1-GFP* is expressed in glia after injury**

(A) Fluorescent microscopic images of uninjured and injured wing nerves 3 dpi. *vir-1-GFP* signal colocalizes with *repo-Gal4* driven myr::tdTomato$^+$ glia after injury. Single optical slices are shown in each case. The arrows show the reporters expressed in glia, while the arrowheads denote the epithelium surrounding the veins. Scare bar: 10 µm. (B) Transcript levels of different *vir-1* isoforms in wings without injury and 2 days after injury. *vir-1 AB* indicates *RA* and *RB* transcript isoforms, *vir-1 ABCGH* refers to *RA, RB, RC, RG* and *RH* isoforms while *vir-1 E* represents the RE isoform. *vir-1* expression levels are normalized to *RpL32* expression. * p = 0.0199, ns (*vir-1 E*) = 0.2433, ns (*vir-1 ABCGH*) = 0.3230. n = 3, respectively.

**Figure S6. Verification of the Su(var)2-10 antibody and control experiments for Fig. 4D**

(A) Single-slice images of wing nerves showing *10xStat92E-GFP* signal upon glial RNAi. RNAi expression was induced by RU486 for 5-7 days after eclosion prior to injury and lasted for the duration of the experiment. *Atg16* RNAi effect is not abrogated by simultaneously introducing a second UAS construct (*UAS-mtdTomato*) arguing against Gal4 unavailability on *Atg16* RNAi and resulting derepression of *10xStat92E-GFP* in Fig. 4D. Scale bar: 5µm. (B) Quantitative analysis of *10xStat92E-GFP* signal shown in (A). Truncated violin plots are shown with median and quartiles. We used unpaired, two-tailed Mann–Whitney test for the analyses. ** p = 0.0033, ns = 0.4491. n = 9, 13, 11, 12. (C) S2 cell extracts transiently expressing Su(var)2-10-EGFP were immunoblotted for GFP (GFP, magenta) and Su(var)2-10 (Su(v), green) simultaneously. Individual channels and the merged image (Mer) are shown. The signals of the reactive bands completely colocalize indicating recognition of Su(var)2-10 by the Su(var)2-10 antibody. A control extract (*C*) not expressing Su(var)2-10::GFP is displayed for comparison on the left. anti-α-tubulin serves as a loading control. (D) Homozygous *Su(var)2-10*$^{MI03442}$ loss-of-function allele and control (*C*) *w*$^{1118}$ L3 stage larva extracts were immunoblotted for Su(var)2-10 and β-actin. Note the reduced Su(var)2-10 isoform band intensities in the mutant in the >~45 kDa size range. Asterisks indicate bands that are not effected by the MiMIC insertion, most likely non-specific bands in larva.

#### Supplementary Table 1

| Gene of interest | | | baseline signal | | signal after injury | |
|---|---|---|---|---|---|---|
| | | | epithelium | nerve | epithelium | nerve |
| AttacinA | **gene-Gal4; UAS-mCD8::GFP** | X | No | No | No | No |
| | **gene-GFP** | X | No | No | Yes | No |
| Drosocin | **gene-Gal4; UAS-mCD8::GFP** | X | No | No | No | No |
| | **gene-GFP** | X | No | No | No | No |
| Metchnikowin | **gene-Gal4; UAS-mCD8::GFP** | - | - | - | - | - |
| | **gene-GFP** | X | No | No | Yes | No |
| Mmp-1 | **gene-Gal4; UAS-mCD8::GFP** | - | - | - | - | - |
| | **gene-GFP** | X | No | No | No | No |

#### Supplementary Table 2

| | | forward primer | reverse primer |
|---|---|---|---|
| ***vir-1* qPCR primers** | vir-1 AB | CGACATTGATGAGCCCAAGTCCA | GGGTGCGCTGGTGTGAAGAT |
| | vir-1 E | AGAGGTGCCATCATTTCCACAAC | GGGTGCGCTGGTGTGAAGAT |
| | vir-1 ABCGH | GTACCATCACGCCCTCAGCC | AGACGGCGGAAGAGATCATCG |
| ***2xmApple-Atg8a* cloning primers** | Apple1 | gaatacaagaagagaactctgaatagggaattgggaattcATGGTGAGCAAGGGCGAGGA | TTGCTCACCATgcccgagcctcctcctttacccttgtacagctcgtccatgccg |
| | Apple2 | tgtacaagggtaaaggaggaggctcgggcATGGTGAGCAAGGGCGAGGA | ATacctcctccgcggccgcctccaccacttcctttgcccttgtacagctcgtccatgccg |
| | Atg8a | ggcaaaggaagtggtggaggcggccgcggaggaggtATGAAGTTCCAATACAAGGAGGAG | ttccttcacaaagatcctctagaggtaccctcgagttaGCCGTAAACATTCTCATCGGAG |
